# Irrational risk aversion in ants is driven by perceptual mechanisms

**DOI:** 10.1101/620054

**Authors:** Massimo De Agrò, Daniel Grimwade, Tomer J. Czaczkes

## Abstract

Animals must often decide between exploiting safe options or risky options with a chance for large gains. While traditional optimal foraging theories assume rational energy maximisation, they fail to fully describe animal behaviour. A logarithmic rather than linear perception of stimuli may shape preference, causing animals to make suboptimal choices. Budget-based rules have also been used to explain risk-preference, and the relative importance of these theories is debated. Eusocial insects represent a special case of risk sensitivity, as they must often make collective decisions based on resource evaluations from many individuals. Previously, colonies of the ant *Lasius niger* were found to be risk-neutral, but the risk preference of individual foragers was unknown. Here, we tested individual *L. niger* in a risk sensitivity paradigm. Ants were trained to associate a scent with 0.55M sucrose solution and another scent with an equal chance of either 0.1 and 1.0M sucrose. Preference was tested in a Y-maze. Ants were extremely risk averse, with 91% choosing the safe option. Even when the risky option offered on average more sucrose (0.8M) than the fixed option, 75% preferred the latter. Based on the psychophysical Weber-Fechner law, we predicted that logarithmically balanced alternatives (0.3M vs 0.1M/0.9M) would be perceived as having equal value. Our prediction was supported, with ants having no preference for either feeder (53% chose the fixed option). Our results thus strongly support perceptual mechanisms driving risk-aversion in ants, and demonstrate that the behaviour of individual foragers can be a very poor predictor of colony-level behaviour.

## Introduction

Finding a good meal is not easy: the environment provides a broad variety of food sources, but individuals are not necessarily able to explore all of them before committing to one (Mehlhorn et al., 2015). The food sources the organism inspects will often have different attributes, and options can be compared in order to choose the best one. This economic decision process is so crucial for organisms that the ability to compare options is found not only in animals, but even in non-neuronal organisms such as plants and slime-moulds (Dener et al., 2016; Reid et al., 2015, 2016).

Traditionally, organisms were assumed to maximise energetic gains while minimising costs, on the basis that evolution should drive animals to have optimal behavioural strategies. However, the optimal foraging theory framework (Pyke et al., 1977) fails to fully describe behaviour - organisms do not always behave optimally. Extensive examples of violation of optimality in animal species can be found, for example, in the literature about risk sensitivity. We define risk as a situation in which the probabilities associated with an option (e.g. food source) are known, but the exact value of it is not. Conversely, “uncertainty” is when not even the probabilities of the various possible payoffs are known.

### Risk sensitivity theories – the budget rule

Risk sensitivity studies were effectively inaugurated by Caraco et al. (1980). They studied the preference of yellow-eyed juncos for different amount of seeds: one of the two alternatives available to the birds was stable, presenting always the same, medium amount of food (safe feeder), while the other one fluctuated in value, but had the same mean pay-out as the safe feeder (risky feeder). The authors then, based on the preference of the animals, designed a utility function (Becker et al., 1964), computing the perceived value (utility) for each number of seeds for the animals. Yellow-eyed juncos presented a concave utility function (and so were risk averse) when in a high energy budget, whereas their utility function was convex (and so they were risk prone) when in a low energy budget. This behaviour was soon formalized as the Energy Budget Rule (Stephens, 1981). However, a growing body of work on risk sensitivity failed to provide consistent empirical support for the budget rule (Kacelnik and Bateson, 1996; Kacelnik and El Mouden, 2013). For this reason the budget rule has recently been reformulated by Lim et. al. (2015). They argue that the classical budget rule is often misused in its binomial interpretation: animals are either risk prone (when in a low energy budget) or risk averse (when in a high energy budget). However, the optimum risk sensitivity in a given situation lies on a continuum, depending on the remaining energy budget of the animal, even arriving at extreme conditions (very low energy budget and very high energy budget) in which risk indifference arise again. Such a continuous interpretation of the budget rule may accommodate results considered as inconsistent the classical budget rule hypothesis (e.g. Hurly, 2003).

### Risk-sensitivity theories – Scalar Utility Theory

An alternative to prescriptive theories (based on optimality modelling) are descriptive theories, which explain behaviours in terms of proximate mechanisms. If risk sensitivity arises as a side-effect of the neural or cognitive architecture of an animal, or due to evolutionary constraints, one need not attempt to fit this behaviour to fitness benefits. A striking pattern in risk preference studies is that animals are often risk averse when risking amounts, but risk seeking when risking delays (Kacelnik and Bateson, 1996). Animals (and humans) are also generally risk averse for potential gains, but risk prone for potential losses (Kahneman and Tversky, 1979). These patterns are elegantly explained by an understanding of how animals perceive the world, as described by Psychophysics (Gescheider, 1976; Stevens, 2017; Tuzlukov, 2013). Stimulus strength has a logarithmic relationship with perception, as formalized by the Weber-Fechner law (Fechner, 1860). Thus, a constant feeder that always presents 5 seeds and a variable feeders presenting alternatively 1 or 9 seeds have the same average; however, 5 seeds are perceived as 5 times more than 1 on a logarithmic curve, while 9 is not even twice as good as 5. Thus, while the mathematical average, and so the true energetic value, of the variable feeder is the same as the one of the safe feeder, it’s geometric average is lower. On logarithmic distributions, such as the Weber-Fechner law by which animals perceive the world, the median is coincident with the geometrical average, and is the measure that describes the overall perceived value of an option, as it is the middle point between the two alternatives. Based on these insights, Kacelnik & El Mouden (2013) developed Scalar Utility Theory (SUT) to describe risk aversion behaviour. They point out that, based on the Weber-Fechner law, the variance of the memory representation of a food value increases as the value itself increases. For this reason, two options with identical mathematical average (means) but different variances will have different medians, with the more variable option having a lower one (see figure 6 from Kacelnik and El Mouden, 2013 for a complete explanation). However, support for this descriptive theory is also mixed: Lim et al. (2015) argue that SUT has even weaker support than the budget rule, with only 8 of the 35 studies reviewed by Kacelnic & Bateson (1996) finding complete risk aversion when risking potential resource gains. Shafir (2000) argued that it is the strength of risk preference that is driven by perceptual mechanisms, while the direction is driven by budget considerations, and could thus accommodate both risk seeking and risk aversion in a manner consistent with logarithmic perception. However, Shafir’s model can only account for alternatives with the same mean value. Whether risk sensitivity is best understood in terms of adaptation or constraints on perceptual mechanisms is thus still under debate.

### Ants as a model for risk sensitivity

Risk sensitivity has been studied in a great variety of animals (for a review, see Kacelnik and El Mouden, 2013). Among those, nectarivores have received particular scrutiny (Perez and Waddington, 1996; Shafir, 2000). The majority of studies on nectarivores have been carried out on bees. Results have, however, been unclear: bees have been observed to be risk indifferent (Banschbach and Waddington, 1994; Fülöp and Menzel, 2000; Perez and Waddington, 1996), risk averse (Shapiro, 2000; Waddington et al., 1981), to follow the budget rule (Cartar, 1991; Cartar and Dill, 1990), or a mixture of those depending on risk variability (Dunlap et al., 2017; Mayack and Naug, 2011; Shafir, 2000; Shafir et al., 1999). Bees and other eusocial insects represent a special case for risk sensitivity. For eusocial insects with non-reproductive workers, the colony is the main unit of selection and a colony can be considered a superorganisms (Boomsma and Gawne, 2018; Hölldobler and Wilson, 2009). As such, the foraging successes of the individual workers are pooled. This buffers colonies against short-term (negative) fluctuation coming from risky choices made by individual foragers individuals. Colonies can also visit multiple food sources simultaneously, allowing them to more efficiently exploit their environment (Czaczkes et al., 2015a; Devigne and Detrain, 2005). Lastly, many eusocial insects can make collective foraging decisions, using recruitment mechanisms to channel workers towards certain resources in the environment (Detrain and Deneubourg, 2008; Gordon, 2019).

While research on risk preference and collective decision-making is extensive, these have rarely been combined. Collective risk sensitivity has been explicitly studied in ants: Burns et al. (2016) presented colonies of rock ants (*Temnothorax albipennis*) a fixed-quality mediocre nest and a variable quality nest. Ants were allowed to explore (and hence evaluate) each nest and then recruited nestmates, and colonies were found to be risk prone. On the other hand, Hübner & Czaczkes (2017) tested the risk sensitivity of black garden ant (*Lasius niger*) colonies to food values. Each colony was presented with two feeders: a stable one, always presenting the same, medium quality sucrose solution (0.55M), and a variable one, presenting alternatively (changing every 3 minutes) either low or high quality sucrose solution (0.1M – 1.0M). Almost all trials showed a clear collective decision for one of the two feeders (as is expected due to symmetry breaking in ants collective decisions, see Beckers et al., 1990, 1993; Czaczkes et al., 2015b; Price et al., 2016), but overall colonies were risk-indifferent: half the colonies chose the safe feeder, and half chose the risky one. This is surprising, as positive feedback from the initially best food source should have resulted in symmetry breaking and a collective choice for that feeder (Beckers et al., 1993; Czaczkes et al., 2015b; Detrain and Deneubourg, 2008; Price et al., 2016).

This work aimed to explore individual risk preference in individual *Lasius niger* ant foragers. Although their collective behaviour appears to be rational, individual workers may not be (Sasaki and Pratt, 2011). They could be victims to the same perceptual constrains discussed above and be strongly influenced by expectations, causing disappointment for some food alternatives, triggering risk sensitivity.

## Materials and Methods

### Subjects

8 queenless *Lasius niger* colony fragments of around 1000 ants were used in the experiment. Each fragment was collected from a different wild colony on the University of Regensburg campus. Colonies fragments forage, deposit pheromone and learn well (Evison et al., 2008; Oberhauser et al., 2018). Each fragment was housed in a transparent plastic box (30×20×40cm), with a layer of plaster on the bottom. A circular plaster nest, 14cm in diameter and 2 cm thick, was also provided. The colonies were kept at room temperature (21-25 c°) and humidity (45-55%), on 12:12 light:dark cycle.

Each colony was fed 0.5mol sucrose solution *ad libitum*, and deprived of food 4 days prior each test. Water was provided *ad libitum* and was always present.

### Experiment 1 – Risk preference between options of equal absolute value

The aim of this experiment was to assess the preference of individual ants between two food sources which provide, on average, an equal amount of sucrose: one feeder provided a stable moderate value (0.55M sucrose, the ‘safe’ option) and one providing a fluctuating value, either high or low (0.1M or 1.0M, the ‘risky’ option). This was achieved by teaching each individual ant to associate each feeder type (risky or safe) with a different odour, and then testing their preference in a Y-maze. Preliminary tests (see ESM1) and previous work (Czaczkes et al., 2018b, 2018c) shows that *L. niger* foragers learn quickly (within 3 visits to each odour) and reliably to associate odours with feeders of different types.

### Training

To begin each experiment ants were allowed onto a 15cm long, 1cm wide runway, with a drop of sucrose at the end. The first ant to encounter the sucrose was marked with a dot of paint, and all other ants were returned to the nest. The marked ant was allowed to drink to satiety and then return to the nest to unload the collected sugar. She was then allowed to make 7 further training visits to the runway and feeder. In each visit we recorded the number of pheromone depositions performed on the runway towards the feeder and towards the nest after foraging. Over the 8 visits the quality and odour of the feeder was varied systematically so that the ant alternately encountered a moderate quality drop of sucrose solution (0.55M, ‘safe’) scented with one odour, or either a low (0.1M) or high (1.0M)(‘risky’) drop of sucrose scented with another odour. These values are clearly distinguishable by the ants (Wendt et al., 2018) and correspond to moderate, low, and high value food sources for *L. niger* (Detrain and Prieur, 2014). Note that the average of the low and high quality solutions equals that of the moderate quality. The solutions were scented using either rosemary or lemon essential oils (0.05 μl per ml). The runway leading to the feeder was covered with a paper overlay scented identically to the sucrose solution being offered. Overlays were scented by storing them in a sealed box containing cotton soaked in essential oil. Overlays were discarded after each return to the nest, to ensure fresh odour and to prevent a build-up of trail pheromone from occurring.

### Testing

After the 8 training visits, the runway was replaced with a Y-maze (arm length 10cm, bifurcation angle 120°). The stem of the Y-maze was overlaid with unscented paper, whereas the two other arms were covered with scented overlays - one bearing the ‘risky’ associated scent, and the other the ‘safe’ associated scent. The maze tapered at the bifurcation to ensure that the ant perceives both scented arms at the same time (following Czaczkes, 2018a). No sucrose was present on the Y-maze. We recorded the ants’ initial arm decision, defined by the ants’ antennae crossing a line 2cm from the bifurcation point. We also recorded the ants’ final decision, defined by the ant crossing a line 8cm from the bifurcation point. However, the initial and final decisions of the ants were almost always the same, and analysis of either choice provides the same results (see ESM1). For brevity we henceforth discuss only the initial decision data. On reaching the end of an arm the ant was allowed to walk onto a piece of paper and brought back to the start of the Y-maze stem, to be retested. The Y-maze test was thus repeated 3 times, to assess reliability of the ant choice. However, this handling may have caused some disruption (see ESM1) and repeated unrewarded trials affect motivation, so we conservatively analysed only the first Y-maze test. After testing, the ant was permanently removed from the colony. In total we tested 64 ants equally divided among 4 different colonies.

For each tested ant, one odour corresponded to the ‘risky’ feeder and one to the ‘safe’ feeder. The association between odour and feeder type, the initial feeder type encountered, the initial value of the ‘risky’ feeder, the side on which the ‘risky’ or ‘safe’ associated odours were presented on the Y-maze test, and the scents associated with the ‘risky’ and ‘safe’ options were all balanced between ants. Performing treatments blind was attempted, but due to the clear negative contrast effects shown by ants on encountering a low quality food source after better ones (Wendt et al., 2018), true blinding was not possible.

### Experiment 2 – Risk preference between options of different absolute value

Experiment 1 demonstrated very strong risk aversion in individual ant foragers. Experiment 2 was designed to test whether risk aversion would be maintained ‘irrationally’, that is, when the ‘risky’ feeder had a higher average quality than the ‘safe’ feeder.

As in experiment 1, the ‘safe’ feeder always presented a medium quality drop (0.55M, indistinguishable for the ants from the solution provided ad libitum to the colony). However, the ‘risky’ feeder alternated between a low quality reward (0.1M) and a very high quality reward (1.5M). The average molarity of the risky feeder (0.8M) was thus higher than the average molarity of the safe one. *L. niger* foragers can distinguish between the three presented molarities (Wendt et al., 2018). Moreover, in a pilot experiment, we observed that when presented with three different molarities ants do learn all three molarities and their associated odours (see ESM1). Each ant was tested on the Y-maze 5 times, but as in experiment 1, only data from the first test was ultimately used (see ESM1). In total we tested 64 ants from 8 new colonies. Each condition (scent association, feeder order, risky feeder order, scent side on the Y-maze) was balanced and equally distributed among colonies.

### Experiment 3 – Risk preference between psychophysically balanced options

One hypothesis explaining the widespread risk aversion found in animals towards reward quantities arises from the psychophysics of perception: intensity is generally perceived logarithmically (Kacelnik and Bateson, 1996; Kacelnik and El Mouden, 2013; see introduction). It is thus the geometrical average between the two risky alternatives that may describe the perceived value. This hypothesis predicts that animals should be indifferent between a safe and a risky option, if the risky option balances the logarithmic differences between the low and high quality reward. In experiment 2, these were not balanced: the geometrical average of the risky feeder (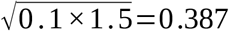) was still lower than the one of the safe feeder 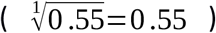, thus the risky option may still have been perceived as worse than the safe option. In this experiment we set out to offer a ‘risky’ option in which the *perceived* qualities of the low and high reward were balanced relative to the moderate reward. We chose a moderate reward of 0.3M, and a low and high reward of 0.1M and 0.9M respectively. The geometrical average of the risky option (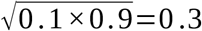) was now equal to the one of the safe option. We thus hypothesised that ants would be indifferent between these two options. Each ant was tested on the Y-maze 5 times, but again only data from the first test was used (see S1). In total we tested 40 ants from 10 different colonies. Each condition (scent association, feeder order, risky feeder order, scent side in the Y-maze) was balanced and equally distributed among colonies.

### Statistical analysis

Statistical analyses were carried out in R 3.3.3 (R Core Team, 2017). Following Forstmeier and Schielzeth (2011), we included in the models only factors and interactions for which we had *a priori* reasons for including. We employed generalized linear mixed effect models using the package lme4 (Bates et al., 2015), with colonies as a random effect. Y-maze choice data was

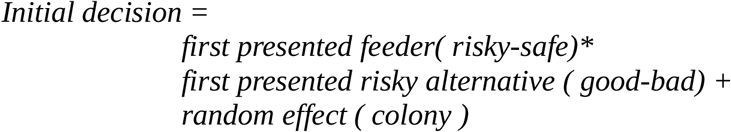

modelled using a binomial distribution and logit link function. We used the following model:

We then used the package car (Fox and Weisberg, 2011) to test which factors of the model had a significant effect on the dependent variable. Subsequently, we carried out post-hoc analysis with Bonferroni correction using the package emmeans (Lenth, 2018) both for the general preference of the ants for either the safe or the risky feeder (safe choice probability against random probability), and for the factors with a significant effect to analyse the direction of the difference. Plots were generated using the package ggplot2 (Wickham, 2009).

Pheromone deposition count was modelled using a poisson distribution and logit link function. Good model fit was confirmed using the DHARMa package (Hartig, 2018), and the pscl package (Jackman, 2017; Zeileis et al., 2008) was used to produce the zero-inflated poisson models when needed. Pheromone deposition was not the focus of the current study, but we include it as descriptive data since it may shed light on how individual perception can shape group choice. We modelled pheromone deposited towards the nest and pheromone deposited on the way back separately, since these are conceptually very different: depositions towards the food reflect the ants’ expectation, and depositions on the return to the nest reflect the ants’ perception. The models used were the following:

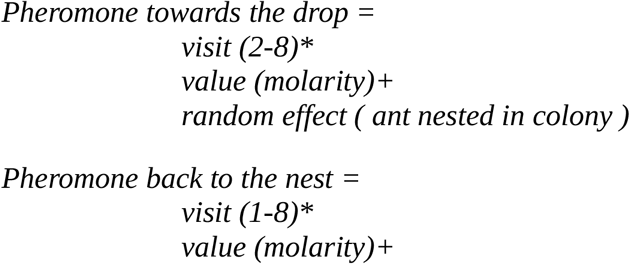

Pheromone deposition data from each of the three experiments were analysed separately, as they were taken by three separate experimenters, and so could not reliably be compared between experiments. Path choice decisions allow much less observer error, so Y-maze data can be pooled between experiments.

Only main results are reported below. For the full analysis see ESM2. The raw data for all the experiments can be found in the supplemental materials ESM3.

## Results

### Experiment 1 – Risk preference between options of equal absolute value

#### Y-maze choice tests

Ants were strongly risk averse, with 91% (58/64) ants initially choosing the safe option (figure 1) (GLMM post-hoc with estimated means, probability=0.911, SE=0.36, z=5.142, p<0.0001). We found no effect of the first presented feeder (GLMM Analysis of Deviance, Chi square=0.709, DF =1, p=0.3), nor of the first presented risky alternative (Chi square=0, DF=1, p=1), nor of the interaction between those two factors (Chi square=0, DF=1, p=1).

**Figure 1:**
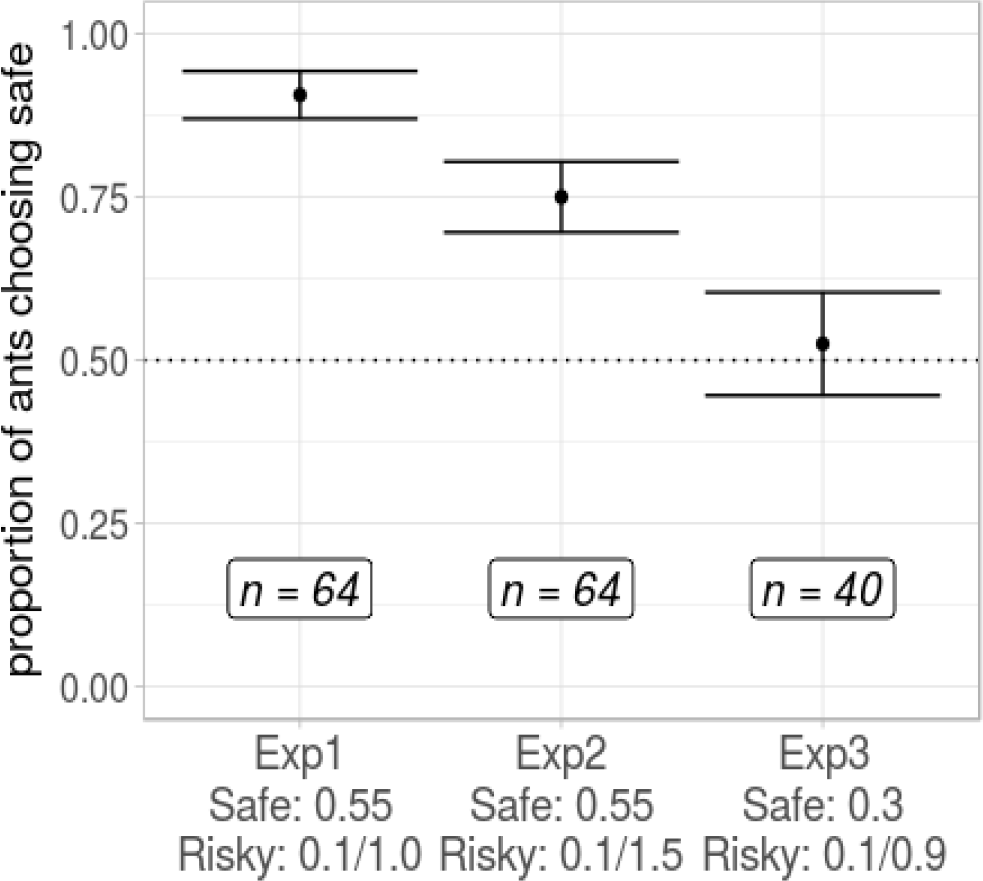
Proportion of ants choosing the safe feeder. Ants preference is different from chance level in experiment 1 (prob.=0.911, SE=0.36, z ratio=5.142, p-value<0.0001) and in experiment 2 (prob.=0.792, SE= 0.068, z ratio = 3.248, p-value =0.001), but not in experiment 3 (prob.=0.535, SE=0.086, z ratio=0.403, p-value=0.687).

#### Pheromone deposition

Considering pheromone deposition towards the feeder, we found an effect of molarity (GLMM Analysis of Deviance, Chi square=12.992, DF=2, p=0.001) and an effect of the interaction between molarity and visit number (GLMM Analysis of Deviance, Chi square=14.469, DF=2, p=0.0007). Specifically, we found that the ants deposited overall more pheromone when going towards the 0.55M drop in comparison to the 1.0M drop (figure 2A, GLMM post-hoc with estimated means, estimate=0.657, SE=0.227, z=2.891, p=0.015). Note that the ant may be expecting to find the 0.1M drop when going towards the 1.0M, because it last experienced the low value associated with that scent. We found no differences in pheromone deposition between the other molarities. Overall, the ants deposited more pheromone on the way to the safe feeder relative to the risky one (GLMM post-hoc with estimated means, estimate=0.498, SE=0.19, z=2.616, p=0.036).

**Figure 2:**
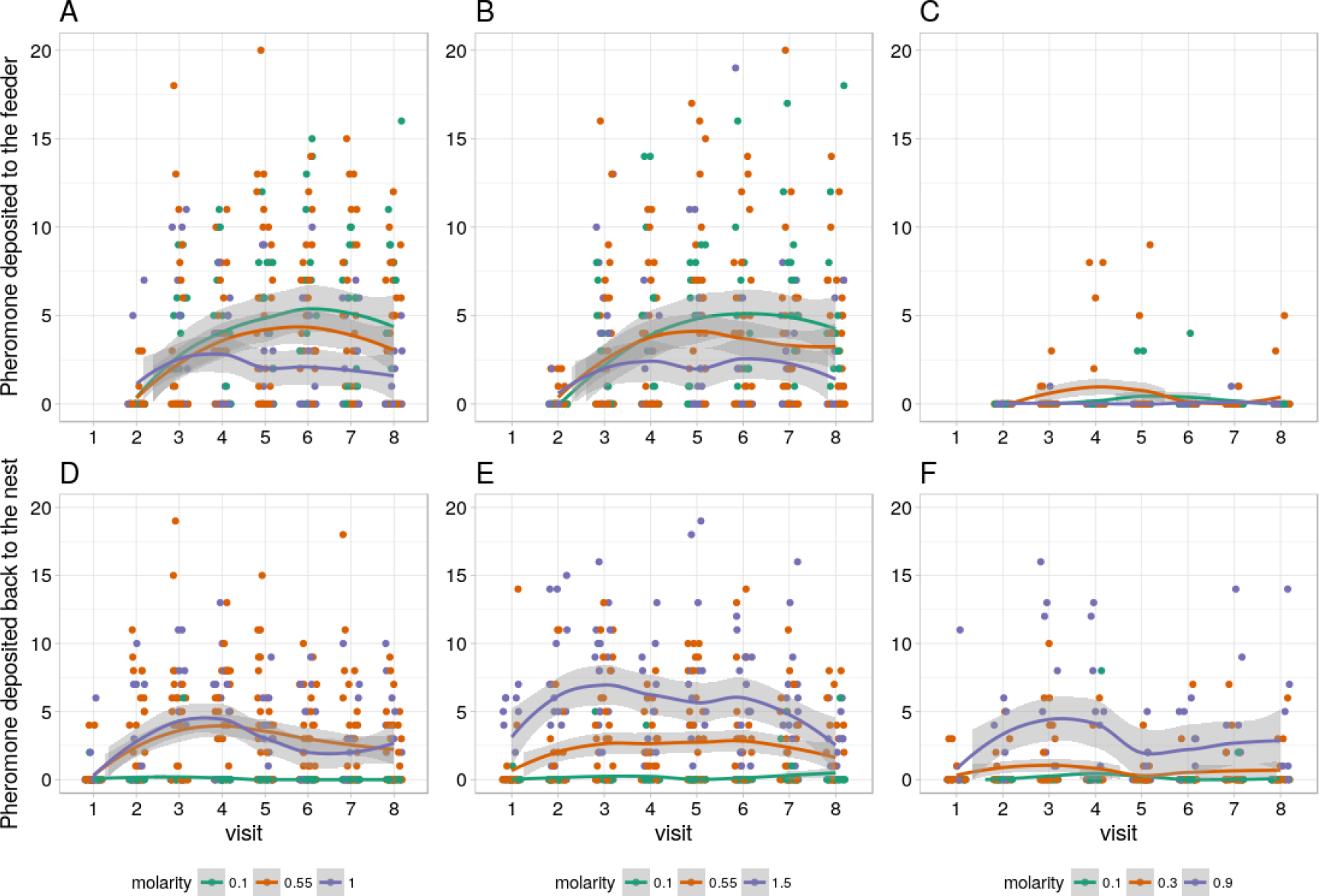
Amount of pheromone deposited by the ants going to the drop and back to the nest across visits in the three experiments. Considering the pheromone deposited on the way to the drop, we found a higher deposition rate for the safe feeder in experiment 1 (A) and in experiment 3 (C) but not in experiment 2 (B). Considering the pheromone deposited on the way back to the nest, we found a higher deposition rate for the safe alternative in experiment 1 (D) and experiment 2 (E), but not in experiment 3 (F).

Considering pheromone deposited when returning to the nest, we found an effect of molarity (GLMM Analysis of Deviance, Chi square=85.97, DF=2, p<0.0001), an effect of visit (GLMM Analysis of Deviance, Chi square=5.11, DF=1, p=0.024), but no effect of their interaction. Specifically, we found that the ants deposited overall less pheromone when going back from the 0.1M drop in comparison to the 0.55M drop (figure 2D, GLMM post-hoc with estimated means, estimate=−2.67, SE=0.154, z=−17.352, p<0.0001) and from the 0.1M drop in comparison to the 1.0M drop (GLMM post-hoc with estimated means, estimate=−2.78, SE=0.194, z=−14.308, p<0.0001). However, there was no difference between the 0.55M drop and the 1.0M drop. Overall the ants deposited more pheromone on the way back from the safe feeder relative to the risky one (GLMM post-hoc with estimated means, estimate=1.28, SE=0.14, z=9.149, p<0.0001).

### Experiment 2 – Risk preference between options of different absolute value

#### Y-maze choice tests

Ants were again strongly risk averse, with 75% (48 / 64) ants initially choosing the safe option (figure 1)(GLMM post-hoc with estimated means, probability=0.792, SE=0.068, z=3.248, p=0.001). We found no effect of the first presented feeder (GLMM Analysis of Deviance, Chi square=2.015, DF=1, p=0.156), nor of the first presented risky alternative (Chi square=0.197, DF=1, p=0.657), nor of the interaction between those two factors (Chi square=1.807, DF=1, p=0.179).

#### Pheromone deposition

The data for the pheromone deposition are summarized in figure 2B and 2E.

Considering pheromone deposited towards the drop, we found an effect of the molarity (figure 2B, GLMM Analysis of Deviance, Chi square=7.489, DF=2, p=0.024). However, post-hoc analysis revealed no difference between any of the molarities: the differences were probably so small that bonferroni correction in the post-hoc analysis brought them above significance.

Considering the pheromone deposited back to the nest, we found an effect of molarity (GLMM Analysis of Deviance, Chi square=133.424, DF=1, p<0.0001), an effect of visit (GLMM, Chi square=10.249, DF=1, p=0.001), and an effect of their interaction (GLMM, Chi square=11.339, DF=2, p=0.003). Ants deposited less pheromone for the 0.1M drop in comparison to the 0.55M drop (figure 2E, GLMM post-hoc with estimated means, estimate=−2.683, SE=0.17, z=−15.742, p<0.0001), less pheromone for the 0.1M in comparison to the 1.5M (GLMM post-hoc with estimated means, estimate=−3.474, SE=0.204, z=−17, p<0.0001) and less for the 0.55M in comparison to the 1.5M (GLMM post-hoc with estimated means, estimate=−0.79, SE=0.19, z=−4.144, p=0.0001). Overall the ants deposited more pheromone on the way back from the safe feeder relative to the risky one (GLMM post-hoc with estimated means, estimate=0.946, SE=0.14, z=6.341, p<0.0001).

### Experiment 3 – Risk preference between psychophysically-balanced options

#### Y-maze choice tests

53% (21/40) of ants chose the safe option (figure 1), a proportion not different from chance (GLMM post-hoc with estimated means, probability=0.535, SE= 0.086, z=0.403, p=0.687).

We found an effect of the first presented feeder (GLMM Analysis of Deviance, Chi square=4.424, DF=1, p=0.0354). Specifically, 71% of the ants that were presented with the safe feeder in visit 1 choose the safe smell during testing, while 35% of the ones presented with the risky feeder first did.

#### Pheromone deposition

Considering pheromone depositions towards the feeder, we found an effect of molarity (GLMM, Chi square=16.133, DF=2, p=0.0003). Ants deposited more pheromone when going towards the 0.3M drop in comparison to the 0.9M drop (figure 2C, GLMM post-hoc with estimated means, estimate=10.444, SE=1.751, z=3.769, p=0.0007), while we found no difference between 0.1M and 0.3M (GLMM post-hoc with estimated means, estimate=0.477, SE=0.174, z=−2.032, p=0.169) and between 0.1M and 0.9M (GLMM post-hoc with estimated means, estimate=4.981, SE=3.452, z=2.317, p=0.082). Overall, ants deposited more pheromone for the safe feeder (GLMM post-hoc with estimated means, estimate=4.679, SE=1.751, z=4.124, p=0.0001)

Considering pheromone deposition back to the nest, we found an effect of molarity (GLMM, Chi square=47.083, DF=2, p<0.0001). Ants deposited less pheromone when returning from the 0.1M drop in comparison to the 0.3M one (figure 2F, GLMM post-hoc with estimated means, estimate=−882, SE=0.143, z=−6144, p<0.0001), less for the 0.1M in comparison to the 0.9M (GLMM post-hoc with estimated means, estimate=−1.479, SE=0.18, z=−8193, p<0.0001) and less for the 0.3M in comparison to the 0.9M (GLMM post-hoc with estimated means, estimate=−0.597, SE=0.165, z=−2.615, p=0.001). Overall the ants deposited the same amount of pheromone on the way back from the safe feeder relative to the risky one (GLMM post-hoc with estimated means, estimate=0.142, SE=0.126, z=1.134, p=1).

## Discussion

Ants show strong risk aversion given equal average payoffs between the risky and safe options (0.1/1.0M vs. 0.55M, experiment 1). Even if the risky option offers 45% higher mean payoffs than the safe reward (0.1M/1.5M vs. 0.55M), ants still show strong risk aversion (experiment 2). We predicted, based on psychophysical principles, that logarithmically-balanced rewards should be perceived as having equal value. We tested this in a situation where the risky reward offered 66% higher payoffs than the safe reward (0.1/0.9M vs 0.3M) and observed, as predicted, indifference between the two options.

### Support for the perceptual basis of risk sensitivity

Our demonstration of risk aversion in resource amounts strongly support the perceptual, descriptive theory of risk sensitivity proposed by Kacelnik & Bateson (1996) and developed by Kacelnik & El Mouden (2013). Specifically, our data suggest functional risk aversion arising from risk neutrality filtered through logarithmic perception. Budget Rule theories (Stephens, 1981) would also predict risk aversion in our context, since the ants are on a positive energy budget – *Lasius niger* would survive for over a week without feeding. However, our ability to accurately predict an indifference point based on logarithmic perception strongly implies that perceptual mechanisms are driving risk aversion in this species. Alternatively, we may have by chance chosen the precise point where logarithmic balancing matches the balance point between improved average gains from a risky option and the premium garnered by a safe bet according to the budget rule. However, this seems unlikely.

The ants in our experiments never showed a preference for the risky alternative. This may seem to imply that the ants were failing to learn the risky option, and associate it with an odour. However, this hypothesis can be ruled out, as it cannot account for the results of experiment 3, where neither food sources is preferred. If the ants were unable to learn the risky option, the only other explanation for experiment 3 would be that a 0.3M is not preferred over complete uncertainty. This can be ruled out, however, as ants clearly prefer 0.3M over 0.1M (ESM1).

### The Budget Rule is neither supported nor refuted

Budget Rule theories (Stephens, 1981) would also predict risk aversion in our context, since the ants are on a positive energy budget – *Lasius niger* would survive for over a week without feeding. However, our ability to accurately predict an indifference point based on logarithmic perception strongly implies that perceptual mechanisms are driving risk aversion in this species. Our data neither supports nor refutes the Budget Rule (Caraco et al., 1980; Lim et al., 2015; Stephens, 1981): we tested all ants after exactly 4 days of starvation, so we cannot know how ants would have behaved on a different energy budget. Lim et al. (2015) strongly critiques SUT, since it predicts suboptimal behaviour, which should be selected against. Logarithmic perception, however, is a widespread phenomenon in the animal kingdom, from roundworms (Luo et al., 2008) to humans (Fechner, 1860), and is argued that the logarithmic scale is the best possible neural representation of magnitudes among other biologically feasible scales (Portugal and Svaiter, 2011). A more precise food evaluation may require more energy than the energy gained from the additional precision. However, this has never been tested in the context of risk sensitivity (Lim et al., 2015). Even if the benefits accrued from a more linear perception of value would outweigh their costs, developmental constraints or pleiotropy may prevent such perception from evolving.

### Lack of support for Prospect Theory

Other theories of risk sensitivity based on perceptual mechanisms exist. Prospect Theory (Kahneman and Tversky, 1979), a hugely influential economic theory of decision-making under risk in humans, predicts that an individual should be risk averse in the context of gains but risk prone in the context of losses. This again derives from logarithmic perception of cumulative gains and losses. However, in Prospect Theory the dividing point between gains and losses is not necessarily at zero. Rather, gains and losses are defined relative to a reference point, which is usually the expected payoff, but may be socially induced (e.g. by comparing ones own salary to that of ones colleagues). Anything above the reference point is perceived as a gain and anything below the reference point is a loss. Disappointment for a lower value after a reference has been established has already been demonstrated in the honeybee (Couvillon and Bitterman, 1984) and ants (Wendt et al., 2018), and suggested in bumblebees (Wiegmann et al., 2003). The reference point for our colonies might have been 0.5M: the solution that the ants are regularly fed on. If this were the case, in experiment 1 the true choice would be between an always neutral value (0.55M, safe), and a risk between a gain (1.0M) and a loss (0.1M). This hypothesis is also supported by the fact that almost no pheromone was deposited for the 0.1M drop. In this case Prospect Theory would still predict risk aversion, as losses are assumed to be perceived more strongly than gains. To test this hypothesis we repeated experiment 1, but with colonies that had been fed *ad libitum* 1.5M sucrose 1 month prior testing (data and procedure can be found in ESM1). If the ants were taking their standard feeding solution as a reference point, every presented solution in this experiment should have been perceived as a loss, and so the ants should have showed risk-seeking. However, we observed the same preference that we saw in the main first experiment – strong risk aversion. Either the ants behaviour is poorly described by Prospect Theory, or the normal feeding solution does not set the reference point. Another possibility is that the reference point is not set by the normal feeding solution, as the four-day food deprivation period may erase the ants memory of the feeding solution. Instead, the reference point could be the most common solution in the current context. In experiment 1 this would be 0.55M, maintaining the same situation of one neutral vs. a loss or a gain, and so predicting the same outcome under Prospect Theory. This hypothesis, however, does not fit the result obtained in experiment 3: if the 0.3M would have been taken as a reference, we should still have observed a preference for the safe option. Either Prospect Theory does not well describe the behaviour of ants, or their reference point remains at 0 in every situation, with every reward being a gain: in the domain of gains Prospect Theory predicts simple logarithmic value perception.

### Risk neutrality at the colony level

Does our understanding of individual behaviour in a risk-choice situation help explain the risk indifference of ants at a colony levels (Hübner and Czaczkes, 2017)? Pheromone deposition rates of individual foragers vary hugely between individuals, even when presented with identical food sources. This is to be expected, given the fact that individual variability may aid collective decisions (Dussutour et al., 2009; O’Shea-Wheller et al., 2017). However, the appropriate measure of pheromone for colony-level decisions is total pheromone deposited. Examining the mean deposition rates for both feeders in experiment 1, we see that ants, on average, deposited more pheromone to the safe feeder (5.5 dots per ant) than the risky feeder (3.9 dots per ants). In Hübner & Czaczkes (2017) each ant made only one or two visits to the feeder, but even when considering only the first two visits ants made more pheromone depositions to the safe (1.5 dots per ant) than to the risky (0.89 dots per ant) feeder. The finding of risk neutrality at the colony level is thus still a puzzle. However, the two experiments are not directly comparable. Firstly, in the current experiment pheromone was removed from the trail every visit. Pheromone presence is known to reduce further pheromone deposition (Czaczkes et al., 2013), perhaps damping out the differences between the two feeders. Secondly, the presence of odours on a path affects pheromone deposition: while pheromone deposition on odourless paths is usually higher on the nestward journey (Beckers et al., 1993; Czaczkes and Heinze, 2015; Czaczkes et al., 2013, 2016), pheromone deposition is higher on outward journeys on scented paths (this study, Czaczkes et al., 2018b, 2018c). Finally, it should be noted that perception of pheromone, much like perception of quality, is also logarithmic (von Thienen et al., 2014), thus emphasising initial differences in pheromone concentration but damping out differences between strong trails. Nevertheless, it seems that colony-level decision-making effectively filters out the ants individual perceptual constrains (this study, Sasaki and Pratt, 2011), but the mechanism used to achieve this is still unknown.

In this study, we found that ants demonstrate risk aversion due to a logarithmic perception of food value. Individual risk preference does not predict colony behaviour, which seems able to filter out perceptual biases.

## Supporting information

Supplementary experiments

Full R script

Raw data

