## Supplementary experiments for "Irrational risk aversion in ants is driven by perceptual mechanisms"

### Supplemental Material 1

### 1 Comparison between initial and final decision & comparison between subsequent testing visits

#### 1.1 Initial and final decision difference

In all three experiments we recorded the ants' initial decision, being the first of the two arms of the Y-maze the ant walked on for at least 2cm. We also recorded the final decision, being the first of the two arms of the Y-maze the ant walked on for at least 8cm. This was done to test the ants' choice robustness. We tested whether the number of ants choosing either arm was different when considering the first decision line or the last decision line, with the following model:

*Safe choice (all tests) =*
*Decision line+*
*random effect ( individual ant nested in colony )*

We found no difference between initial and final decision in any of the experiment (table S1). This means that the ants' preference is robust and they rarely change Y maze arm after having made their initial choice. We thus decided to focus further analysis only on initial choice.

| | Odds ratio | Standard Error | Z ratio | p-value |
|---|---|---|---|---|
| Experiment 1 | 0.814 | 0.234 | -0.715 | 0.4746 |
| Experiment 2 | 1.1 | 0.195 | 0.539 | 0.59 |
| Experiment 3 | 1.103 | 0.243 | 0.445 | 0.656 |

*Table S1 – results of the post-hoc analysis for the three experiment checking for difference between initial and final decision.*

#### 1.2 Subsequent testing visits

In all three experiment after training we repeatedly tested the trained ants (3 times for the first experiment, 5 times for the other two). This was also done to assess the ants choice robustness. To assess if there was a difference between the subsequent testing visits we used the following model:

*Primary safe choice (all tests) =*
*Testing visit (1-3/1-5)+*
*random effect ( individual ant nested in colony )*

We found that in both experiment 1 and experiment 2 the percentage of ants choosing the safe alternative decreased over subsequent visits (Table S2). This was most likely due to the ant stopping their attempts to choose the preferred feeder after not finding the reward in the first or second testing visit, with many individuals starting instead to perform random search. Some ants may also have been disrupted by the handling. We thus decided to include only the first choice each ant made in the final analysis, as we felt that this better represents the ants' true preference. The percentage of ants choosing safe did not change over subsequent visits in experiment 3 (table S2). However, this was to be expected, as the ants were already choosing at chance level, so no reversion to random search could be detected. For correctness and consistency with the previous experiments, we chose to focus our analysis on only the first initial decision of each ant. Our analysis was thus identical to that used in experiments 1 and 2.

| | Estimate | Standard Error | Z ratio | p-value |
|---|---|---|---|---|
| Experiment 1 | -0.762 | 0.2551 | -3.109 | 0.0004 |
| Experiment 2 | -0.20978 | 0.086 | -2.426 | 0.015 |
| Experiment 3 | -0.07576 | 0.104 | -0.727 | 0.467 |

*Table S2 – effect of the factor "Testing visit" on the percentage of ant choosing safe in the three experiments. Note that for experiment 1 we had to drop "colony" as a random factor, because the model could not converge otherwise.*

### 2 Control experiments

#### 2.1 Ant perception of 0.1, 0.3 and 0.9 sucrose molarities

In experiment 3, the ants were presented with feeders offering 0.1, 0.3 and 0.9 molar sucrose. Relative to experiments 1 and 2, the medium and the low quality drop had very similar molarities in absolute terms, so that we decided to run a pilot experiment to test whether the ant could discriminate, and subsequently choose reliably between, the three molarities.

We ran two testing blocks. In the first, the ants were presented with two drops of different molarities, 0.1 and 0.3, and were trained to associate each to a smell. We followed the methodology described for the main experiment (see methods section in the main paper), alternating the presentation of the low quality alternative and the high quality alternative in the 8 training visits. Afterwards we tested the ants in the Y-maze, repeating the test 5 times. The second experiment was identical to the first, but the ants were presented with 0.3 and 0.9 molarities. We did this last block just as a control, since as 0.3 and 0.9 are further apart than 0.55 and 1.0 we were confident that they could discriminate between the two. We tested 20 ants for each block (40 in total), stemming from 6 different colonies. First, we tested the robustness of the ants' choices, checking whether with subsequent visits the number of ants choosing the high value drop decreased. We modelled as follows:

*High value choice (all tests) =*
*Testing visit (1-5)+*
*Contrast (0.1vs0.3 or 0.3vs0.9)*
*random effect ( individual ant nested in colony )*

We found that the ants did not change their preference over subsequent visits for either of the two contrasts (table S3). This could be because this task is easier than the risk vs safe evaluation, having to compare only two molarities in which one is definitely better than the other. For the subsequent analysis, we kept all 5 testing visits. We modelled the data as follows:

Then, we ran a post-hoc test to check which of the groups differed from chance level. We found that the ants significantly preferred 0.3 over 0.1 when considering both the first decision line and the second decision line. However, we found that the ants did not significantly preferred 0.9 over 0.3, remaining at chance level (Table S4). This was surprising to us, as the contrast between 0.9 and 0.3 should be easier to sense, or at least equally difficult if the ants follow a logarithmic perception, and in both cases easier than the contrast between 0.55 and 1.0. We suspect that, due to the lower sample size in these experiments, we have experienced a type II error (false negative). However, we decided to present our data as it is, without a post-hoc increase in sample size, following good scientific practice.

*High value choice (all tests) =*
*Decision line+*
*Contrast (0.1vs0.3 or 0.3vs0.9)*
*random effect ( individual ant nested in colony )*

| Factor | Chi-square | Degrees of freedom | p-value |
|---|---|---|---|
| Contrast | 3.5857 | 1 | 0.058 |
| Testing visit | 1.46 | 1 | 0.227 |
| Contrast:Testing visit | 0.024 | 1 | 0.876 |

*Table S3 – Analysis of deviance (Type II chi-square test) of the model to check difference between testing visit. For this model we had to drop colony as random factor because the model did not converge otherwise. Note that both Testing visit and the interaction between Testing visit and the contrast are not significant. We can conclude that there is no difference between the test visits and there is no difference between the two contrast in testing visits change.*

| Contrast | Decision line | probability | Standard Error | Z ratio | p-value |
|---|---|---|---|---|---|
| 0.1 vs 0.3 | First decision line | 0.86 | 0.057 | 3.844 | 0.0005 |
| 0.1 vs 0.3 | Last decision line | 0.87 | 0.054 | 3.988 | 0.0003 |
| 0.3 vs 0.9 | First decision line | 0.648 | 0.095 | 1.461 | 0.576 |
| 0.3 vs 0.9 | Last decision line | 0.724 | 0.085 | 2.264 | 0.094 |

*Table S4 – post-hoc analysis of the probability of ants choosing the high value alternative, bonferroni corrected.*

#### 2.2 Ant preference among 3 molarities

In the main experiment the ants were presented with three different food qualities, and were required to remember all three in order to make a choice between the two feeders. We decided to run a pilot experiment on order to test whether the ants could remember three molarities, rather than just the best one among others.

The ants performed 9 sequential visits to a runway, identical to the one of the main experiment. At the end of the runway the ant may find either a 1.5M drop, always unscented, a 1.0M drop, either rosemary or lemon scented, and a 0.25M drop, scented with the other odour (see table S5).

| Visit 1 | Visit 2 | Visit 3 | Visit 4 | Visit 5 | Visit 6 | Visit 7 | Visit 8 | Visit 9 |
|---|---|---|---|---|---|---|---|---|
| 1.5M | **1.0M** | **0.25M** | 1.5M | **1.0M** | **0.25M** | 1.5M | **1.0M** | **0.25M** |
| 1.5M | **0.25M** | **1.0M** | 1.5M | **0.25M** | **1.0M** | 1.5M | **0.25M** | **1.0M** |

*Table S5 – Training visit sequence for the three molarity experiment. Bold text represent scented visits. The same molarity always have the same scent.*

Afterwards we tested the ant preference between the 1.0M scent and the 0.25M scent in the Y-maze, repeating the test 5 times. If the ants could only learn the best alternative among the presented ones, they should choose randomly between the second best and the worst. However, if the ants can remember and compare all three values, they should prefer the 1.0M.

We planned to test 32 ants coming from 8 different colonies. However, after having tested 15 ants coming from 5 different colonies we decided to stop the pilot, given the clear preference of the animals: On the first trial, both initial and final decision, 100% of the ants choose the scent associated with 1.0M. We observed a decrease on the ants performance in subsequent files, however the overall percentage remained at 92%. While we are aware that stopping an experiment prematurely when results are as expected can lead to type I errors (false positives), we felt that the unambiguous nature of these results warranted doing so here.

#### 3 Risk preference in the context of losses (maintained on 1.5M sucrose)

Prospect Theory predicts that individuals should be risk averse in the context of gains and risk prone in the context of losses. The reference point from which we decide if something is a gain or a loss is not necessarily 0: we may take an expected value as a reference. For ants, this value it could be the feeding solution they are maintained on, normally 0.5M. We decided to replicate experiment 1 (see main paper) with 4 colonies that had been fed *ad libitum* 1.5M sucrose instead of the usual 0.5M for one month prior testing. 63 ants were tested in total. Training and testing procedure were identical to those described in the main paper. We found that 82% (52/63) of the ants preferred the safe alternative. All data and analysis are provided in supplement S2
