## Supplementary material for "Irrational risk aversion in ants is driven by perceptual mechanisms": Full R script

### Supplemental material 2

#### Data Analysis

**Intro**

This supplement provides the entire R script and output of the statistical analysis we performed and figures produced, in their original form. It is presented in the spirit of open and transparent science, but has not been carefully curated.

### 1 Column descriptions

| column_name | description |
|---|---|
| date | Testing date |
| colony | Colony number |
| antN | Ant number |
| condition | Experimental condition |
| antID | Ant individual ID |
| safescent | Scent of the safe feeder |
| riskscent | Scent of the risky feeder |
| firstrisk | Which of the risky alternatives is presented first |
| firstfeed | Which of the two feeders is presented first |
| visit | Visit number |
| type | Training or testing visit |
| mol | molarity of the drop in the current visit |
| lastmol | molarity of the drop in the previous visit |
| lastfeedmol | molarity of the drop in the previous visit of the same feeder |
| scent | scent of the current visit |
| feed | feeder of the current visit |
| phergo | pheromone deposited on the way to the drop |
| pherbk | pheromone deposited on the way back to the nest |
| safeside | side of the safe smell in the Y maze test |
| firstchoice | initial choice side |
| endchoice | final choice side |
| firstchoicesafe | initial choice binomial data (safe is 1) |
| endchoicesafe | final choice binomial data (safe is 1) |

### 2 Data analysis

first I load packages

```
library(lme4)
library(DHARMa)
library(car)
library(emmeans)
library(reshape2)
library(ggplot2)
library(knitr)
library(pscl)

set.seed(123)#set seed for replicability in random simulations
```

#### 2.1 Binomial Choice

##### 2.1.1 Preliminary questions

###### 2.1.1.1 initial vs. final

first, I want to know if initial and final choice differ

###### 2.1.1.1.1 exp 1

```
fsdiff<-melt(risksa, measure.vars = c("firstchoicesafe","endchoicesafe"))

mdiff<-glmer(value~variable+(1|colony/antID),data=fsdiff,family="binomial")
Anova(mdiff)
```

```
## Analysis of Deviance Table (Type II Wald chisquare tests)
##
## Response: value
##           Chisq Df Pr(>Chisq)
## variable 0.5111  1     0.4746
```

```
e<-emmeans(mdiff, ~variable, type="response")
pairs(e)
```

```
##  contrast                        odds.ratio        SE  df z.ratio p.value
##  firstchoicesafe / endchoicesafe  0.8144338 0.2338279 Inf  -0.715  0.4746
##
## Tests are performed on the log odds ratio scale
```

there is no difference between initial and final choice, I will now on only use the initial for further analysis

###### 2.1.1.1.2 exp 2

```
fsdiff<-melt(riskirr, measure.vars = c("firstchoicesafe","endchoicesafe"))

mdiff<-glmer(value~variable+(1|colony/antID),data=fsdiff,family="binomial")
Anova(mdiff)
```

```
## Analysis of Deviance Table (Type II Wald chisquare tests)
##
## Response: value
##           Chisq Df Pr(>Chisq)
## variable 0.2903  1       0.59
```

```
e<-emmeans(mdiff, ~variable, type="response")
pairs(e)
```

```
##  contrast                        odds.ratio        SE  df z.ratio p.value
##  firstchoicesafe / endchoicesafe   1.100189 0.1949616 Inf   0.539  0.5900
##
## Tests are performed on the log odds ratio scale
```

there is no difference between initial and final choice, I will now on only use the initial for further analysis

###### 2.1.1.1.3 exp 3

```
fsdiff<-melt(riskgeo, measure.vars = c("firstchoicesafe","endchoicesafe"))

mdiff<-glmer(value~variable+(1|colony/antID),data=fsdiff,family="binomial")
Anova(mdiff)
```

```
## Analysis of Deviance Table (Type II Wald chisquare tests)
##
## Response: value
##           Chisq Df Pr(>Chisq)
## variable 0.1981  1     0.6563
```

```r
e<-emmeans(mdiff, ~variable, type="response")
pairs(e)
```

```
##  contrast                        odds.ratio        SE  df z.ratio p.value
##  firstchoicesafe / endchoicesafe   1.102893 0.2426951 Inf   0.445  0.6563
##
## Tests are performed on the log odds ratio scale
```

there is no difference between initial and final choice, I will now on only use the initial for further analysis

###### 2.1.1.2 vistits n.

now, I want to know if the visits differ from one another

###### 2.1.1.2.1 exp 1

```r
risksa$visit<-as.numeric(risksa$visit)
mvisdiff<-glmer(firstchoicesafe~visit+(1|colony/antID),data=risksa,family="binomial",
                glmerControl(optimizer="bobyqa", optCtrl = list(maxfun = 1000000000)))
```

```
## Warning in checkConv(attr(opt, "derivs"), opt$par, ctrl = control
## $checkConv, : Model failed to converge with max|grad| = 0.0157403 (tol =
## 0.001, component 1)
```

```r
mvisdiff<-glmer(firstchoicesafe~visit+(1|antID),data=risksa,family="binomial",
                glmerControl(optimizer="bobyqa", optCtrl = list(maxfun = 1000000000)))
Anova(mvisdiff)
```

```
## Analysis of Deviance Table (Type II Wald chisquare tests)
##
## Response: firstchoicesafe
##       Chisq Df Pr(>Chisq)
## visit 9.668  1   0.001875 **
## ---
## Signif. codes:  0 '***' 0.001 '**' 0.01 '*' 0.05 '.' 0.1 ' ' 1
```

```r
summary(mvisdiff)
```

```
## Generalized linear mixed model fit by maximum likelihood (Laplace
##   Approximation) [glmerMod]
##  Family: binomial  ( logit )
## Formula: firstchoicesafe ~ visit + (1 | antID)
##    Data: risksa
## Control:
## glmerControl(optimizer = "bobyqa", optCtrl = list(maxfun = 1e+09))
##
##      AIC      BIC   logLik deviance df.resid
##    196.7    206.4    -95.3    190.7      189
##
## Scaled residuals:
##     Min      1Q  Median      3Q     Max
## -2.8804  0.3022  0.3472  0.5081  0.8520
##
## Random effects:
##  Groups Name        Variance Std.Dev.
##  antID  (Intercept) 0.3172   0.5632
```

```
## Number of obs: 192, groups:  antID, 64
##
## Fixed effects:
##             Estimate Std. Error z value Pr(>|z|)
## (Intercept)   9.0752     2.5512   3.557 0.000375 ***
## visit        -0.7616     0.2449  -3.109 0.001875 **
## ---
## Signif. codes:  0 '***' 0.001 '**' 0.01 '*' 0.05 '.' 0.1 ' ' 1
##
## Correlation of Fixed Effects:
##       (Intr)
## visit -0.996
```

the percentage of ants choosing safe decreases with successive visits. this means that more and more ants after not finding the sugar drop start doing a random search. I will from now on only observe the first visit, being it a clearer indication of choice

###### 2.1.1.2.2 exp 2

```r
mvisdiff<-glmer(firstchoicesafe~visit+(1|colony/antID),data=riskirr,family="binomial",
                glmerControl(optimizer="bobyqa", optCtrl = list(maxfun = 100000)))
Anova(mvisdiff)
```

```
## Analysis of Deviance Table (Type II Wald chisquare tests)
##
## Response: firstchoicesafe
##        Chisq Df Pr(>Chisq)
## visit 5.8851  1    0.01527 *
## ---
## Signif. codes:  0 '***' 0.001 '**' 0.01 '*' 0.05 '.' 0.1 ' ' 1
```

```r
summary(mvisdiff)
```

```
## Generalized linear mixed model fit by maximum likelihood (Laplace
##   Approximation) [glmerMod]
##  Family: binomial  ( logit )
## Formula: firstchoicesafe ~ visit + (1 | colony/antID)
##    Data: riskirr
## Control:
## glmerControl(optimizer = "bobyqa", optCtrl = list(maxfun = 1e+05))
##
##      AIC      BIC   logLik deviance df.resid
##    415.4    430.4   -203.7    407.4      316
##
## Scaled residuals:
##     Min      1Q  Median      3Q     Max
## -1.8473 -1.1416  0.6020  0.7331  1.0362
##
## Random effects:
##  Groups       Name        Variance Std.Dev.
##  antID:colony (Intercept) 0.14277  0.3778
##  colony       (Intercept) 0.03706  0.1925
## Number of obs: 320, groups:  antID:colony, 64; colony, 8
##
## Fixed effects:
```

```
##             Estimate Std. Error z value Pr(>|z|)
## (Intercept)  2.96566    0.97581   3.039  0.00237 **
## visit       -0.20978    0.08647  -2.426  0.01527 *
## ---
## Signif. codes:  0 '***' 0.001 '**' 0.01 '*' 0.05 '.' 0.1 ' ' 1
##
## Correlation of Fixed Effects:
##       (Intr)
## visit -0.988
```

the percentage of ants choosing safe decreases with successive visits. this means that more and more ants after not finding the sugar drop start doing a random search. I will from now on only observe the first visit, being it a clearer indication of choice

###### 2.1.1.2.3 exp 3

```r
mvisdiff<-glmer(firstchoicesafe~visit+(1|colony/antID),data=riskgeo,family="binomial",
                glmerControl(optimizer="bobyqa", optCtrl = list(maxfun = 100000)))
Anova(mvisdiff)
```

```
## Analysis of Deviance Table (Type II Wald chisquare tests)
##
## Response: firstchoicesafe
##        Chisq Df Pr(>Chisq)
## visit 0.5282  1     0.4674
```

```r
summary(mvisdiff)
```

```
## Generalized linear mixed model fit by maximum likelihood (Laplace
##   Approximation) [glmerMod]
##  Family: binomial  ( logit )
## Formula: firstchoicesafe ~ visit + (1 | colony/antID)
##    Data: riskgeo
## Control:
## glmerControl(optimizer = "bobyqa", optCtrl = list(maxfun = 1e+05))
##
##      AIC      BIC   logLik deviance df.resid
##    282.4    295.6   -137.2    274.4      196
##
## Scaled residuals:
##     Min      1Q  Median      3Q     Max
## -1.1864 -0.9476  0.6974  0.9092  1.2280
##
## Random effects:
##  Groups       Name        Variance Std.Dev.
##  antID:colony (Intercept) 0.3117   0.5583
##  colony       (Intercept) 0.0000   0.0000
## Number of obs: 200, groups:  antID:colony, 40; colony, 10
##
## Fixed effects:
##             Estimate Std. Error z value Pr(>|z|)
## (Intercept)  0.94158    1.16051   0.811    0.417
## visit       -0.07576    0.10425  -0.727    0.467
##
## Correlation of Fixed Effects:
```

```
##        (Intr)
## visit -0.989
```

there is no difference between visits. I will use only first for consistency, but I expect random choice. in this case, it is clear why there is no decrease: if the choice is already random there is no room for reverting to random choice with subsequent visits.

##### 2.1.2 modeling

now to the actual model. I drop antID because I kept only one observation for each ant

###### 2.1.2.1 Exp 1

```r
risksasing<-subset(risksa,risksa$visit==9)

mExp1<-glmer(firstchoicesafe~firstfeed*firstrisk+(1|colony),data=risksasing,family="binomial",
             glmerControl(optimizer="bobyqa", optCtrl = list(maxfun = 100000)))

simres<-simulateResiduals(mExp1) #standard seed for random values is 123
plot(simres, asFactor=T)
```

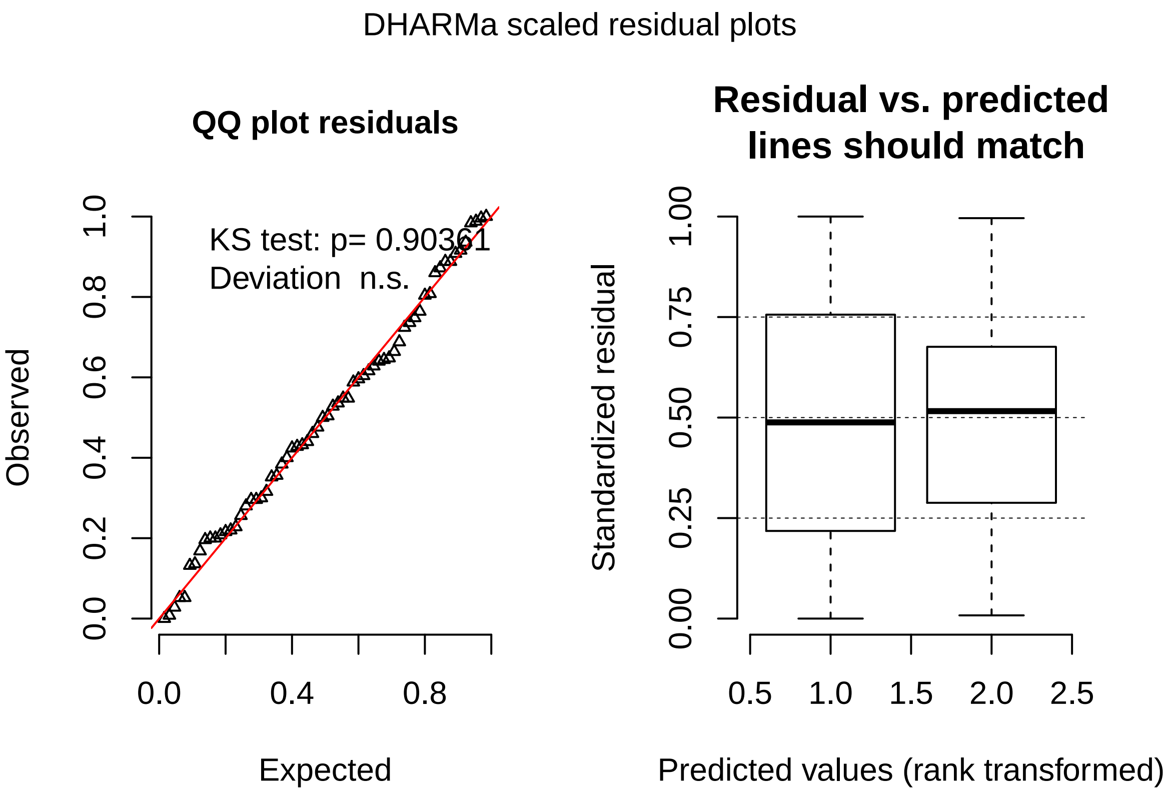

model is good here

```r
Anova(mExp1)
```

```
## Analysis of Deviance Table (Type II Wald chisquare tests)
##
## Response: firstchoicesafe
##                       Chisq Df Pr(>Chisq)
## firstfeed            0.7092  1     0.3997
```

```
## firstrisk           0.0000  1      1.0000
## firstfeed:firstrisk 0.0000  1      1.0000
```

no effect of any of the factors. will just test overall preference

```
meanobj <- emmeans(mExp1,~1, type="response")
kable(test(meanobj))
```

| 1 | prob | SE | df | z.ratio | p.value |
|---|---|---|---|---|---|
| overall | 0.911087 | 0.0366564 | Inf | 5.142427 | 3e-07 |

ants prefer the safe **91%**

###### 2.1.2.2 Exp 2

```
riskirrsing<-subset(riskirr,riskirr$visit==9)

mExp2<-glmer(firstchoicesafe~firstfeed*firstrisk+(1|colony),data=riskirrsing,family="binomial",
             glmerControl(optimizer="bobyqa", optCtrl = list(maxfun = 100000)))

simres<-simulateResiduals(mExp2) #standard seed for random values is 123
plot(simres, asFactor=T)
```

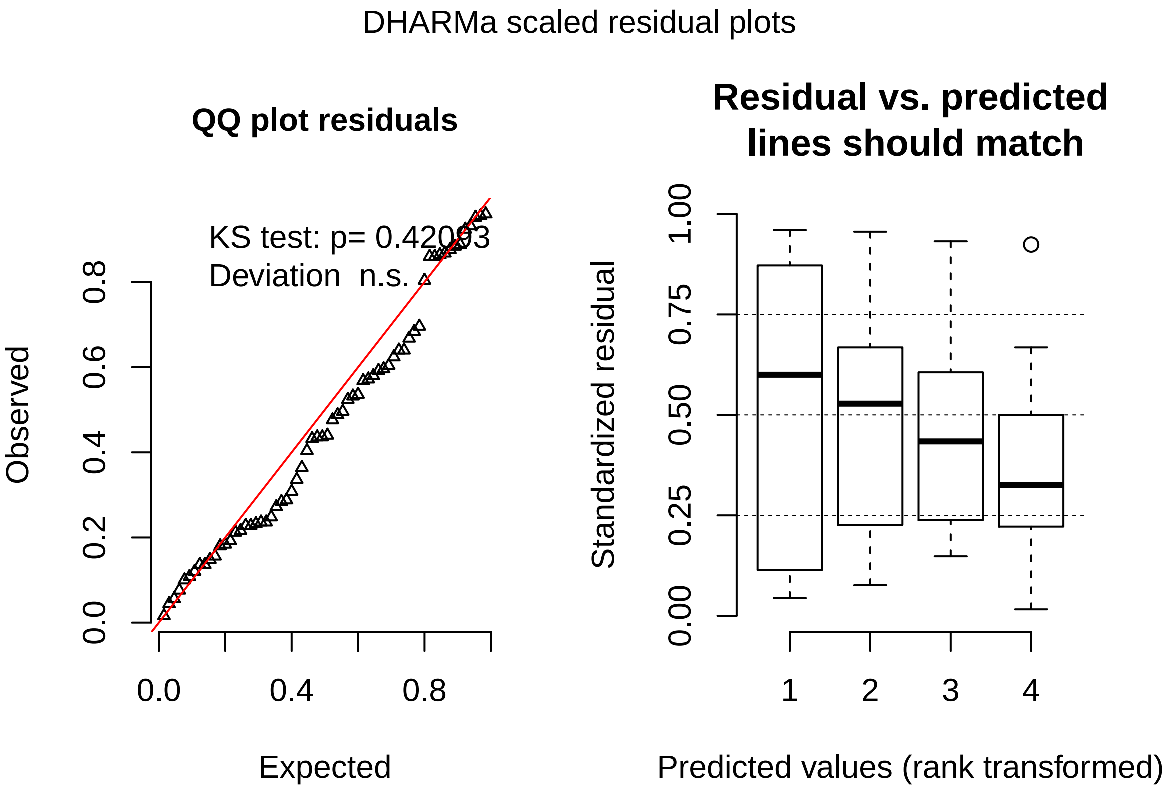

good model also here

```
Anova(mExp2)
```

```
## Analysis of Deviance Table (Type II Wald chisquare tests)
##
```

```
## Response: firstchoicesafe
##                         Chisq Df Pr(>Chisq)
## firstfeed              2.0148  1     0.1558
## firstrisk              0.1969  1     0.6572
## firstfeed:firstrisk 1.8066  1     0.1789
```

still, no effect of factors.

```
meanobj<-emmeans(mExp2,~1, type="response")
kable(test(meanobj))
```

| 1 | prob | SE | df | z.ratio | p.value |
|---|---|---|---|---|---|
| overall | 0.7918708 | 0.0677956 | Inf | 3.248403 | 0.0011605 |

ants prefere the safe **79%**

###### 2.1.2.3 Exp 3

```
riskgeosing<-subset(riskgeo,riskgeo$visit==9)

mExp3<-glmer(firstchoicesafe~firstfeed*firstrisk+(1|colony),data=riskgeosing,family="binomial",
             glmerControl(optimizer="bobyqa", optCtrl = list(maxfun = 100000)))

simres<-simulateResiduals(mExp3) #standard seed for random values is 123
plot(simres, asFactor=T)
```

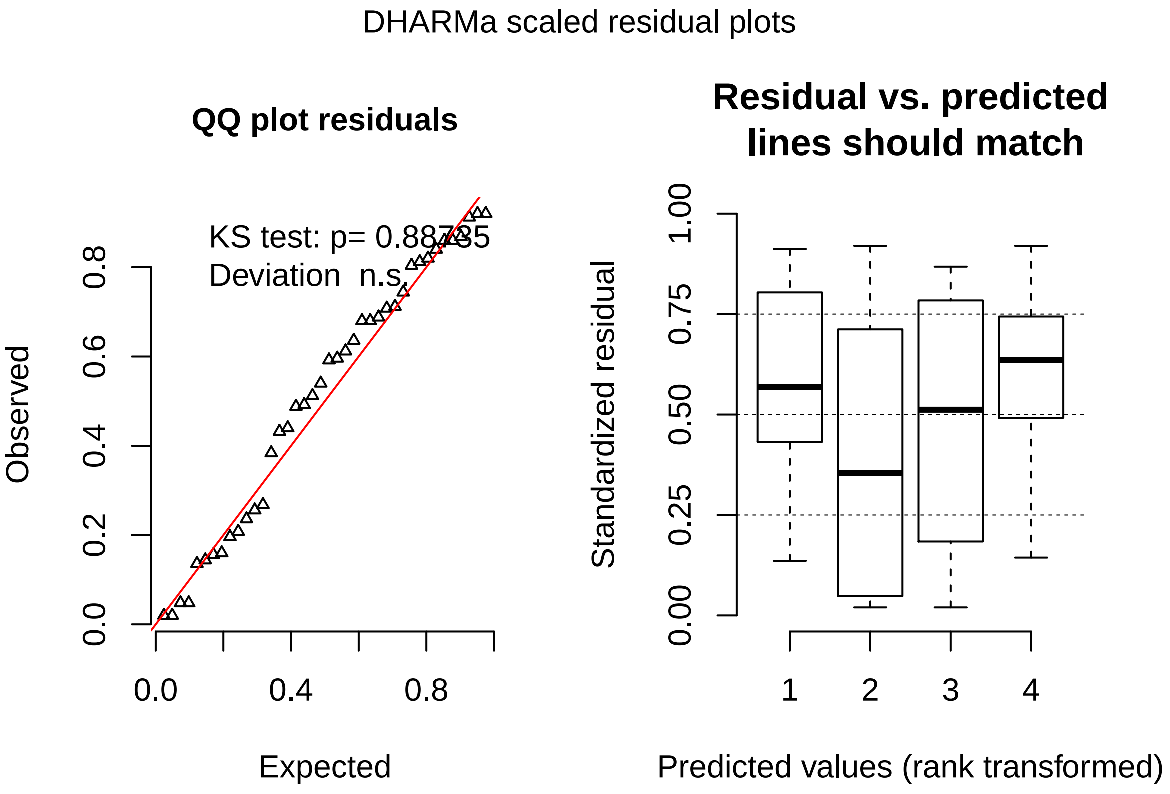

still, good model

```r
Anova(mExp3)
```

```
## Analysis of Deviance Table (Type II Wald chisquare tests)
##
## Response: firstchoicesafe
##                      Chisq Df Pr(>Chisq)
## firstfeed           4.4237  1    0.03544 *
## firstrisk           0.0146  1    0.90388
## firstfeed:firstrisk 0.6679  1    0.41377
## ---
## Signif. codes:  0 '***' 0.001 '**' 0.01 '*' 0.05 '.' 0.1 ' ' 1
```

there is an effect of the first presented feeder. first of all let's look at overall percentage

```r
meanobj<-emmeans(mExp3,~1, type="response")
kable(test(meanobj))
```

| 1 | prob | SE | df | z.ratio | p.value |
|---|---|---|---|---|---|
| overall | 0.534919 | 0.0863723 | Inf | 0.4029695 | 0.6869707 |

ants prefere the safe **53%**.

```r
meanobj<-emmeans(mExp3,~firstfeed, type="response")
```

```
## NOTE: Results may be misleading due to involvement in interactions
```

```r
pairs(meanobj)
```

```
##  contrast     odds.ratio        SE  df z.ratio p.value
##  risky / safe  0.2159797 0.1499687 Inf  -2.207  0.0273
##
## Results are averaged over the levels of: firstrisk
## Tests are performed on the log odds ratio scale
```

```r
meanobj
```

```
##  firstfeed      prob        SE  df asymp.LCL asymp.UCL
##  risky     0.3483315 0.1072458 Inf 0.1747432 0.5743483
##  safe      0.7122198 0.1042953 Inf 0.4772311 0.8702891
##
## Results are averaged over the levels of: firstrisk
## Confidence level used: 0.95
## Intervals are back-transformed from the logit scale
```

more ants go to the safe when this is presented fist, more ants go to the risky when is presented first. overall is random! probably in front of a random choice they just go for the first experienced.

##### 2.1.3 Graph together

need to calculate a full model to get SE in order to plot.

```r
risksing<-subset(risk,risk$visit==9)
library(plyr)
risksing$condition<-revalue(risksing$condition, c("GeomAvrg"="Exp3", "Irrational"="Exp2","sameAvg"="Exp

mTot <- glmer(firstchoicesafe~condition+(1|colony),data=risksing,family="binomial",
```

```
               glmerControl(optimizer="bobyqa", optCtrl = list(maxfun = 100000)))

Anova(mTot)
```

```
## Analysis of Deviance Table (Type II Wald chisquare tests)
##
## Response: firstchoicesafe
##            Chisq Df Pr(>Chisq)
## condition 16.918  2   0.000212 ***
## ---
## Signif. codes:  0 '***' 0.001 '**' 0.01 '*' 0.05 '.' 0.1 ' ' 1
```

```
meanobj<-emmeans(mTot,~condition, type="response")
toplot1<-as.data.frame(meanobj)
kable(pairs(meanobj))
```

| contrast | odds.ratio | SE | df | z.ratio | p.value |
|---|---|---|---|---|---|
| Exp3 / Exp2 | 0.3684211 | 0.1578563 | Inf | -2.330468 | 0.0516981 |
| Exp3 / Exp1 | 0.1143376 | 0.0609494 | Inf | -4.068170 | 0.0001402 |
| Exp2 / Exp1 | 0.3103449 | 0.1604340 | Inf | -2.263396 | 0.0610622 |

```
ggplot(toplot1,aes(x=condition,y=prob))+
  ylab("proportion of ants choosing safe")+
  ylim(0,1)+
  scale_x_discrete(name= NULL,
                   limits=c("Exp1","Exp2","Exp3"),
                   labels=c("Exp1" = "Exp1\nSafe: 0.55\nRisky: 0.1/1.0",
                            "Exp2" = "Exp2\nSafe: 0.55\nRisky: 0.1/1.5",
                            "Exp3" = "Exp3\nSafe: 0.3\nRisky: 0.1/0.9"))+
  theme_light()+
  theme(axis.text.x = element_text(size=12),
        axis.text.y = element_text(size=12),
        axis.title.y = element_text(size=14))+
  geom_label(x=1, y=0.15, label="n = 64",size=5, aes(fontface=3))+
  geom_label(x=2, y=0.15, label="n = 64",size=5, aes(fontface=3))+
  geom_label(x=3, y=0.15, label="n = 40",size=5, aes(fontface=3))+
  geom_hline(yintercept = 0.5,linetype="dotted")+
  geom_point()+
  geom_errorbar(aes(ymin=prob-SE,ymax=prob+SE))
```

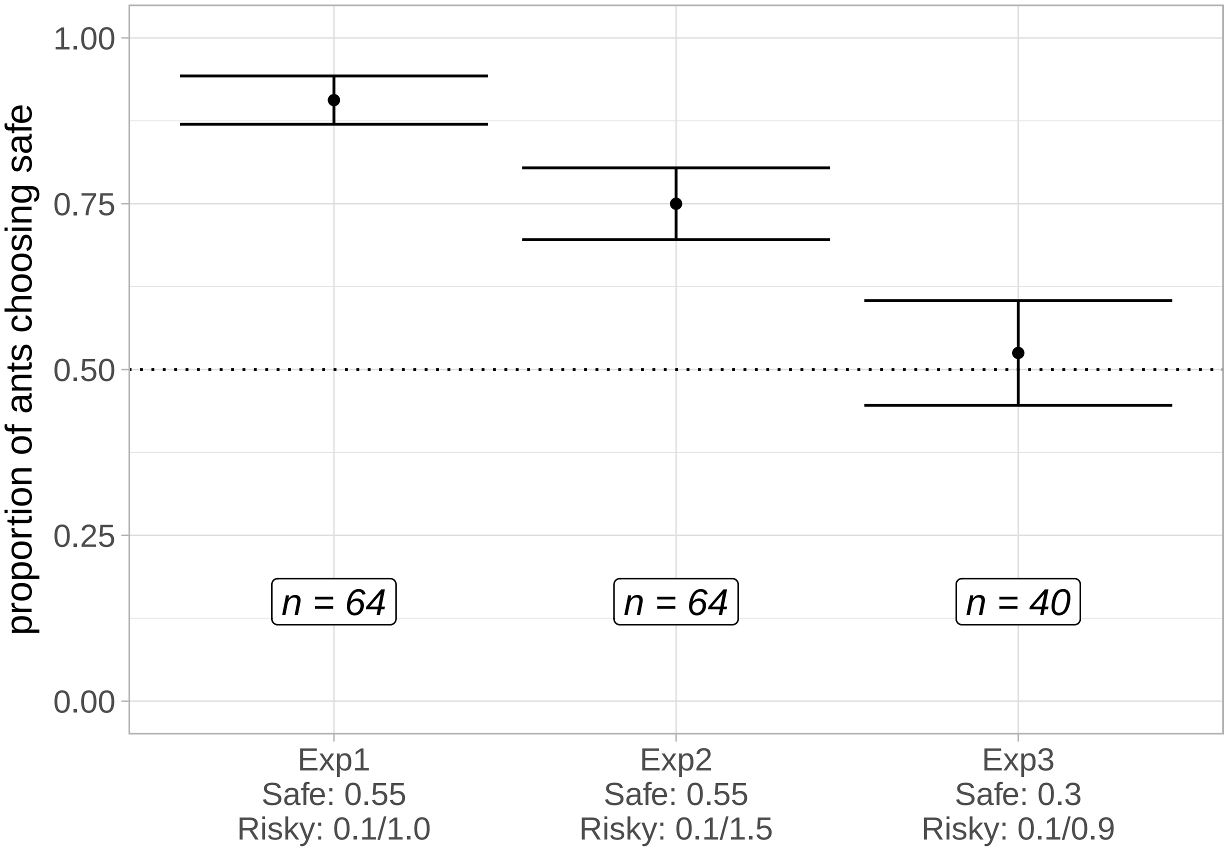

#### 2.2 Pheromone deposition

I will look at the pheromone deposited on the way to the drop and back to the nest for each experiment across visits.

##### 2.2.1 Exp1

###### 2.2.1.1 To the drop

```
risksa$visit<-as.numeric(risksa$visit)

mpExp1<-glmer(phergo~visit*mol+(1|colony/antID),data=risksa,family="poisson",
              glmerControl(optimizer="bobyqa", optCtrl = list(maxfun = 100000)))
simres<-simulateResiduals(mpExp1) #standard seed for random values is 123
plot(simres, asFactor=T)
```

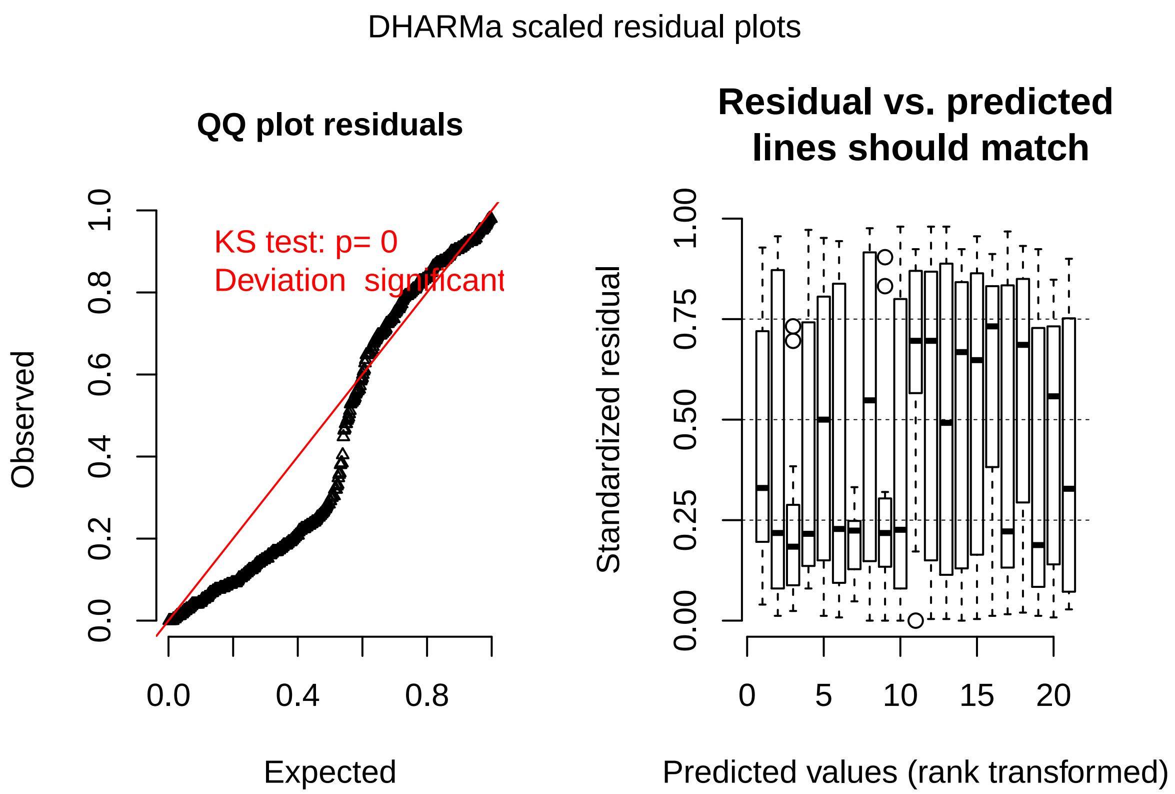

the model is zero inflated. let's remodel

```
mpExp1 <- zeroinfl(phergo ~ visit*mol + 1 | colony/antID, data = risksa)
```

```
## Error in optim(fn = loglikfun, gr = gradfun, par = c(start$count, start$zero, : valore non finito fo
```

does not work. I will remove colony from random effect.

```
mpExp1 <- zeroinfl(phergo ~ visit*mol + 1 | antID, data = risksa)

Anova(mpExp1)
```

```
## Analysis of Deviance Table (Type II tests)
##
## Response: phergo
##           Df   Chisq Pr(>Chisq)
## visit      1  1.7587  0.1847906
## mol        2 12.9922  0.0015093 **
## visit:mol  2 14.4692  0.0007212 ***
## ---
## Signif. codes:  0 '***' 0.001 '**' 0.01 '*' 0.05 '.' 0.1 ' ' 1
```

```
meanobj<-emmeans(mpExp1,~visit*mol,type="response")
contrast(meanobj,list(mol0.1vs0.55=c(1,-1,0),
                      mol0.1vs1.0=c(1,0,-1),
                      mol0.55vs1.0=c(0,1,-1),
                      SafeVsRisky=c(-0.5,1,-0.5)),
         adjust="bonferroni")
```

```
##  contrast       estimate        SE  df z.ratio p.value
##  mol0.1vs0.55 -0.3385364 0.2244369 Inf  -1.508  0.5258
```

```
##  mol0.1vs1.0   0.3189496 0.2433582 Inf   1.311  0.7599
##  mol0.55vs1.0  0.6574860 0.2274213 Inf   2.891  0.0154
##  SafeVsRisky   0.4980112 0.1903691 Inf   2.616  0.0356
##
## Results are averaged over the levels of: antID
## P value adjustment: bonferroni method for 4 tests
```

```
risksap<-subset(risksa,risksa$visit<9) #just remove tests
risksap <- droplevels(risksap)

phg1<-ggplot(risksap,aes(x=visit,y=phergo, color=mol))+
  labs(title = "A")+
  scale_color_brewer(name="molarity", palette="Dark2")+
  ylab("Pheromone deposited to the feeder")+
  scale_x_discrete(limits=c(1,2,3,4,5,6,7,8))+
  ylim(0,20)+
  theme_light()+
  theme(axis.text.x = element_text(size=12,colour="black"),
        axis.text.y = element_text(size=12,colour="black"),
        axis.title.x = element_blank(),
        axis.title.y = element_text(size=14),
        plot.title = element_text(size=18),
        legend.position="none")+
  geom_jitter(width = 0.2,height=0)+
  geom_smooth()
```

###### 2.2.1.2 Back to the nest

```
mpExp1<-glmer(pherbk~visit*mol+(1|colony/antID),data=risksa,family="poisson",
              glmerControl(optimizer="bobyqa", optCtrl = list(maxfun = 100000)))
simres<-simulateResiduals(mpExp1) #standard seed for random values is 123
plot(simres, asFactor=T)
```

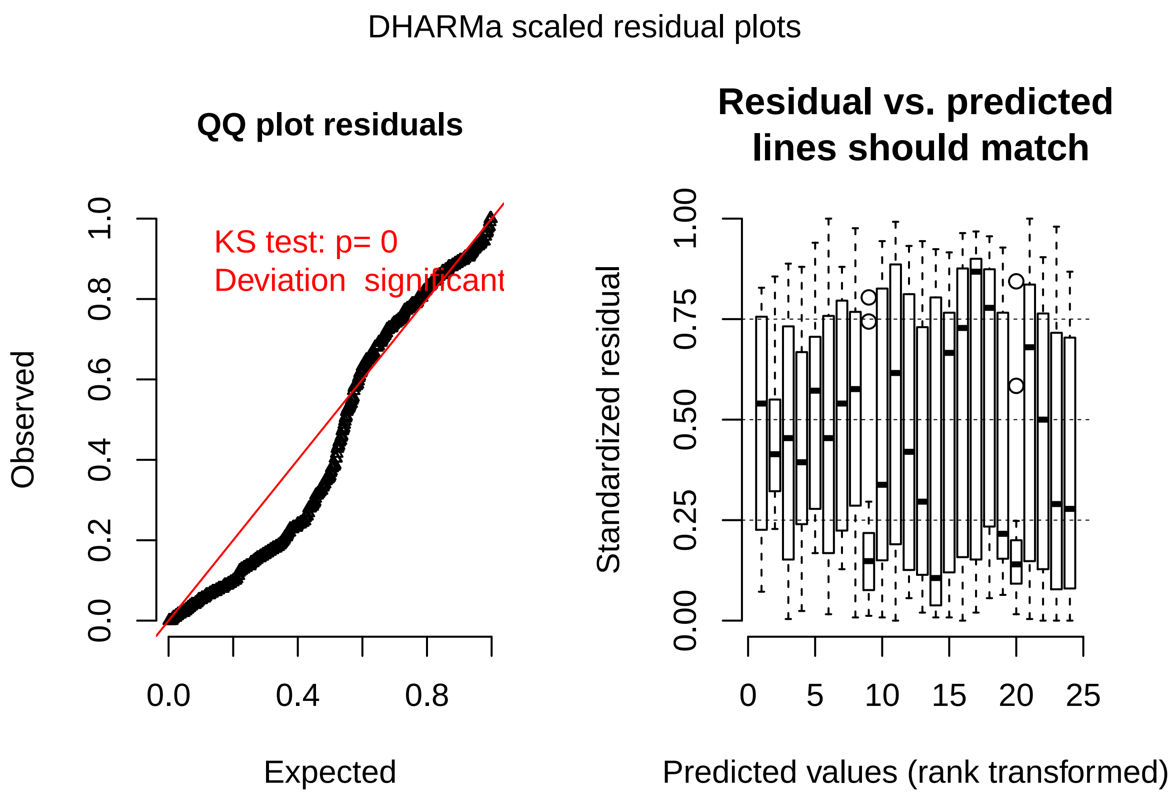

the model is zero inflated. let's remodel

```
mpExp1 <- zeroinfl(pherbk ~ visit*mol + 1 | colony/antID, data = risksa)
```

```
## Error in optim(fn = loglikfun, gr = gradfun, par = c(start$count, start$zero, : valore non finito fo
```

does not work. I will remove colony from random effect.

```
mpExp1 <- zeroinfl(pherbk ~ visit*mol + 1 | antID, data = risksa)

Anova(mpExp1)
```

```
## Analysis of Deviance Table (Type II tests)
##
## Response: pherbk
##           Df   Chisq Pr(>Chisq)
## visit      1  5.1128    0.02375 *
## mol        2 85.9726    < 2e-16 ***
## visit:mol  2  3.9549    0.13842
## ---
## Signif. codes:  0 '***' 0.001 '**' 0.01 '*' 0.05 '.' 0.1 ' ' 1
```

```
meanobj<-emmeans(mpExp1,~visit*mol,type="response")
contrast(meanobj,list(mol0.1vs0.55=c(1,-1,0),
                      mol0.1vs1.0=c(1,0,-1),
                      mol0.55vs1.0=c(0,1,-1),
                      SafeVsRisky=c(-0.5,1,-0.5)),
         adjust="bonferroni")
```

```
##  contrast       estimate        SE  df z.ratio p.value
##  mol0.1vs0.55 -2.6697986 0.1538646 Inf -17.352  <.0001
```

```
##  mol0.1vs1.0  -2.7803883 0.1943198 Inf -14.308  <.0001
##  mol0.55vs1.0 -0.1105897 0.1852878 Inf  -0.597  1.0000
##  SafeVsRisky   1.2796045 0.1398674 Inf   9.149  <.0001
##
## Results are averaged over the levels of: antID
## P value adjustment: bonferroni method for 4 tests
```

```
phb1<-ggplot(risksap,aes(x=visit,y=pherbk, color=mol))+
  labs(title = "D")+
  scale_color_brewer(name="molarity", palette="Dark2")+
  ylab("Pheromone deposited back to the nest")+
  scale_x_discrete(limits=c(1,2,3,4,5,6,7,8))+
  ylim(0,20)+
  theme_light()+
  theme(axis.text.x = element_text(size=12,colour="black"),
        axis.text.y = element_text(size=12,colour="black"),
        axis.title.x = element_text(size=14),
        axis.title.y = element_text(size=14),
        plot.title = element_text(size=18),
        legend.position="bottom")+
  geom_jitter(width = 0.2,height=0)+
  geom_smooth()
```

##### 2.2.2 Exp2

###### 2.2.2.1 To the drop

```
riskirr$visit<-as.numeric(riskirr$visit)

mpExp2<-glmer(phergo~visit*mol+(1|colony/antID),data=riskirr,family="poisson",
              glmerControl(optimizer="bobyqa", optCtrl = list(maxfun = 100000)))
simres<-simulateResiduals(mpExp2) #standard seed for random values is 123
plot(simres, asFactor=T)
```

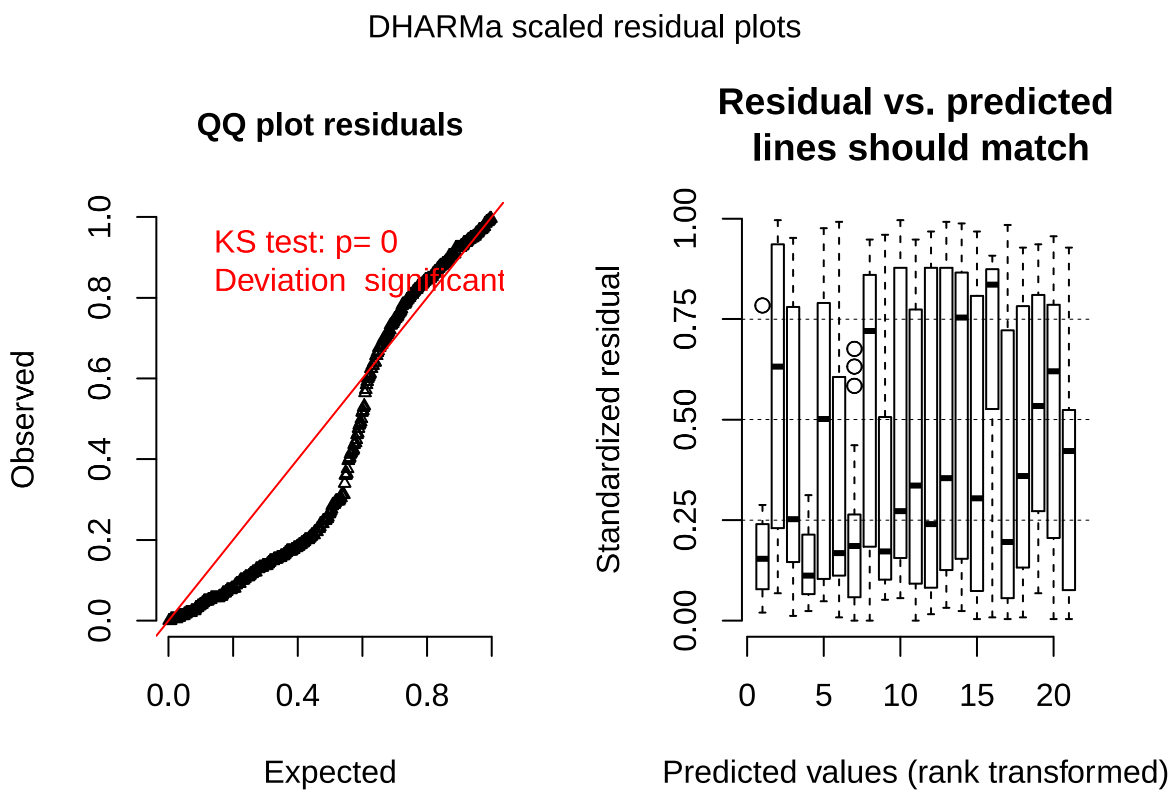

the model is zero inflated. let's remodel

```
mpExp2 <- zeroinfl(phergo ~ visit*mol + 1 | colony/antID, data = riskirr)
```

```
## Error in optim(fn = loglikfun, gr = gradfun, par = c(start$count, start$zero, : valore non finito fo
```

does not work. I will remove colony from random effect.

```
mpExp2 <- zeroinfl(phergo ~ visit*mol + 1 | antID, data = riskirr)

Anova(mpExp2)
```

```
## Analysis of Deviance Table (Type II tests)
##
## Response: phergo
##           Df  Chisq Pr(>Chisq)
## visit      1 0.2798    0.59680
## mol        2 7.4888    0.02365 *
## visit:mol  2 1.6650    0.43495
## ---
## Signif. codes:  0 '***' 0.001 '**' 0.01 '*' 0.05 '.' 0.1 ' ' 1
```

```
meanobj<-emmeans(mpExp2,~visit*mol,type="response")
contrast(meanobj,list(mol0.1vs0.55=c(1,-1,0),
                      mol0.1vs1.5=c(1,0,-1),
                      mol0.55vs1.5=c(0,1,-1),
                      SafeVsRisky=c(-0.5,1,-0.5)),
         adjust="bonferroni")
```

```
##  contrast       estimate        SE  df z.ratio p.value
##  mol0.1vs0.55 -0.1742026 0.2293294 Inf  -0.760  1.0000
```

```
##  mol0.1vs1.5   0.3228436 0.2652927 Inf   1.217  0.8945
##  mol0.55vs1.5  0.4970462 0.2332662 Inf   2.131  0.1324
##  SafeVsRisky   0.3356244 0.1894927 Inf   1.771  0.3061
##
## Results are averaged over the levels of: antID
## P value adjustment: bonferroni method for 4 tests
```

```
riskirrp<-subset(riskirr,riskirr$visit<9) #just remove tests
riskirrp <- droplevels(riskirrp)

phg2<-ggplot(riskirrp,aes(x=visit,y=phergo, color=mol))+
  labs(title = "B")+
  scale_color_brewer(name="molarity", palette="Dark2")+
  ylab("Pheromone deposited to the feeder")+
  scale_x_discrete(limits=c(1,2,3,4,5,6,7,8))+
  ylim(0,20)+
  theme_light()+
  theme(axis.text.x = element_text(size=12,colour="black"),
        axis.text.y = element_text(size=12,colour="black"),
        axis.title.x = element_blank(),
        axis.title.y = element_blank(),
        plot.title = element_text(size=18),
        legend.position="none")+
  geom_jitter(width = 0.2,height=0)+
  geom_smooth()
```

###### 2.2.2.2 Back to the nest

```
mpExp2<-glmer(pherbk~visit*mol+(1|colony/antID),data=riskirr,family="poisson",
              glmerControl(optimizer="bobyqa", optCtrl = list(maxfun = 100000)))
simres<-simulateResiduals(mpExp2) #standard seed for random values is 123
plot(simres, asFactor=T)
```

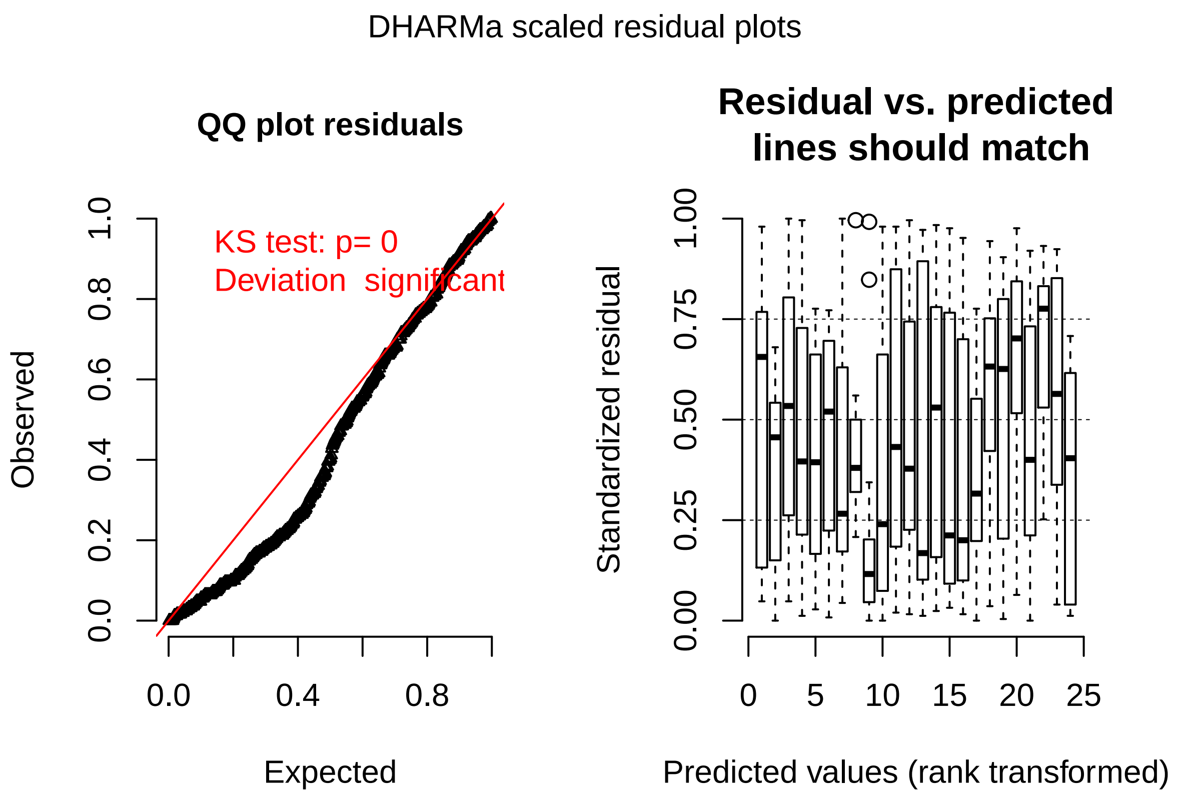

the model is zero inflated. let's remodel

```
mpExp2 <- zeroinfl(pherbk ~ visit*mol + 1 | colony/antID, data = riskirr)
```

```
## Error in optim(fn = loglikfun, gr = gradfun, par = c(start$count, start$zero, : valore non finito fo
```

does not work. I will remove colony from random effect.

```
mpExp2 <- zeroinfl(pherbk ~ visit*mol + 1 | antID, data = riskirr)

Anova(mpExp2)
```

```
## Analysis of Deviance Table (Type II tests)
##
## Response: pherbk
##           Df   Chisq Pr(>Chisq)
## visit      1  10.249   0.001368 **
## mol        2 133.424  < 2.2e-16 ***
## visit:mol  2  11.339   0.003449 **
## ---
## Signif. codes:  0 '***' 0.001 '**' 0.01 '*' 0.05 '.' 0.1 ' ' 1
```

```
meanobj<-emmeans(mpExp2,~visit*mol,type="response")
contrast(meanobj,list(mol0.1vs0.55=c(1,-1,0),
                      mol0.1vs1.5=c(1,0,-1),
                      mol0.55vs1.5=c(0,1,-1),
                      SafeVsRisky=c(-0.5,1,-0.5)),
         adjust="bonferroni")
```

```
##  contrast       estimate        SE  df z.ratio p.value
##  mol0.1vs0.55 -2.6835091 0.1704700 Inf -15.742  <.0001
```

```
##  mol0.1vs1.5  -3.4739039 0.2043446 Inf -17.000  <.0001
##  mol0.55vs1.5 -0.7903948 0.1907377 Inf  -4.144  0.0001
##  SafeVsRisky   0.9465571 0.1492691 Inf   6.341  <.0001
##
## Results are averaged over the levels of: antID
## P value adjustment: bonferroni method for 4 tests
```

```
phb2<-ggplot(riskirrp,aes(x=visit,y=pherbk, color=mol ))+
  labs(title = "E")+
  scale_color_brewer(name="molarity", palette="Dark2")+
  ylab("Pheromone deposited back to the nest")+
  scale_x_discrete(limits=c(1,2,3,4,5,6,7,8))+
  ylim(0,20)+
  theme_light()+
  theme(axis.text.x = element_text(size=12,colour="black"),
        axis.text.y = element_text(size=12,colour="black"),
        axis.title.x = element_text(size=14),
        axis.title.y = element_blank(),
        plot.title = element_text(size=18),
        legend.position="bottom")+
  geom_jitter(width = 0.2,height=0)+
  geom_smooth()
```

##### 2.2.3 Exp3

###### 2.2.3.1 To the drop

```
riskgeo$visit<-as.numeric(riskgeo$visit)

mpExp3<-glmer(phergo~visit*mol+(1|colony/antID),data=riskgeo,family="poisson", glmerControl(optimizer="
simres<-simulateResiduals(mpExp3) #standard seed for random values is 123
plot(simres, asFactor=T)
```

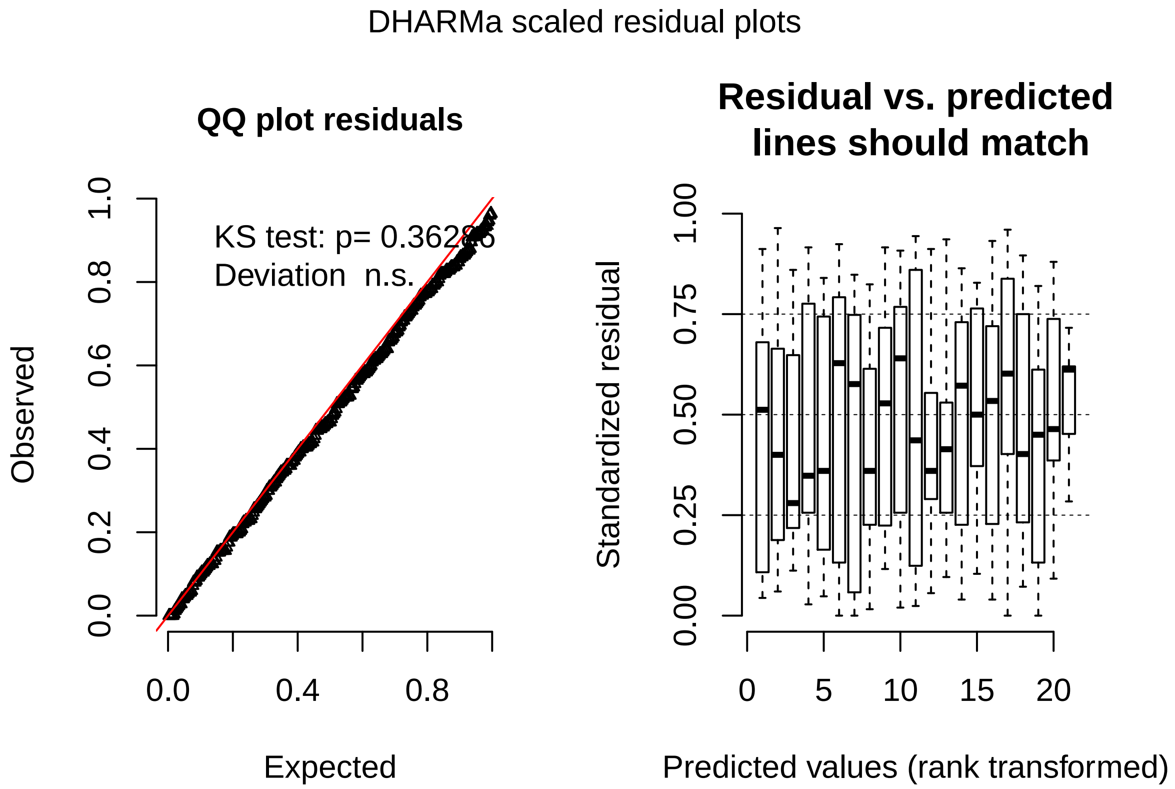

```
Anova(mpExp3)
```

```
## Analysis of Deviance Table (Type II Wald chisquare tests)
##
## Response: phergo
##              Chisq Df Pr(>Chisq)
## visit       0.2874  1  0.5918885
## mol        16.1336  2  0.0003138 ***
## visit:mol   3.7139  2  0.1561452
## ---
## Signif. codes:  0 '***' 0.001 '**' 0.01 '*' 0.05 '.' 0.1 ' ' 1
```

```
meanobj<-emmeans(mpExp3,~visit*mol,type="response")
contrast(meanobj,list(mol0.1vs0.3=c(1,-1,0),
                      mol0.1vs0.9=c(1,0,-1),
                      mol0.3vs0.9=c(0,1,-1),
                      SafeVsRisky=c(-0.5,1,-0.5)),
         adjust="bonferroni")
```

```
##  contrast          ratio        SE  df z.ratio p.value
##  mol0.1vs0.3  0.4769786 0.1737865 Inf  -2.032  0.1687
##  mol0.1vs0.9  4.9814816 3.4526207 Inf   2.317  0.0821
##  mol0.3vs0.9 10.4438260 6.5008615 Inf   3.769  0.0007
##  SafeVsRisky  4.6792943 1.7508890 Inf   4.124  0.0001
##
## P value adjustment: bonferroni method for 4 tests
## Tests are performed on the log scale
```

```
riskgeop<-subset(riskgeo,riskgeo$visit<9) #just remove tests
riskgeop <- droplevels(riskgeop)
```

```r
phg3<-ggplot(riskgeop,aes(x=visit,y=phergo, color=mol))+
  labs(title = "C")+
  scale_color_brewer(name="molarity", palette="Dark2")+
  ylab("Pheromone deposited to the feeder")+
  scale_x_discrete(limits=c(1,2,3,4,5,6,7,8))+
  ylim(0,20)+
  theme_light()+
  theme(axis.text.x = element_text(size=12,colour="black"),
        axis.text.y = element_text(size=12,colour="black"),
        axis.title.x = element_blank(),
        axis.title.y = element_blank(),
        plot.title = element_text(size=18),
        legend.position="none")+
  geom_jitter(width = 0.2,height=0)+
  geom_smooth()
```

###### 2.2.3.2 Back to the nest

```r
mpExp3<-glmer(pherbk~visit*mol+(1|colony/antID),data=riskgeo,family="poisson",
              glmerControl(optimizer="bobyqa", optCtrl = list(maxfun = 100000)))
simres<-simulateResiduals(mpExp3) #standard seed for random values is 123
plot(simres, asFactor=T)
```

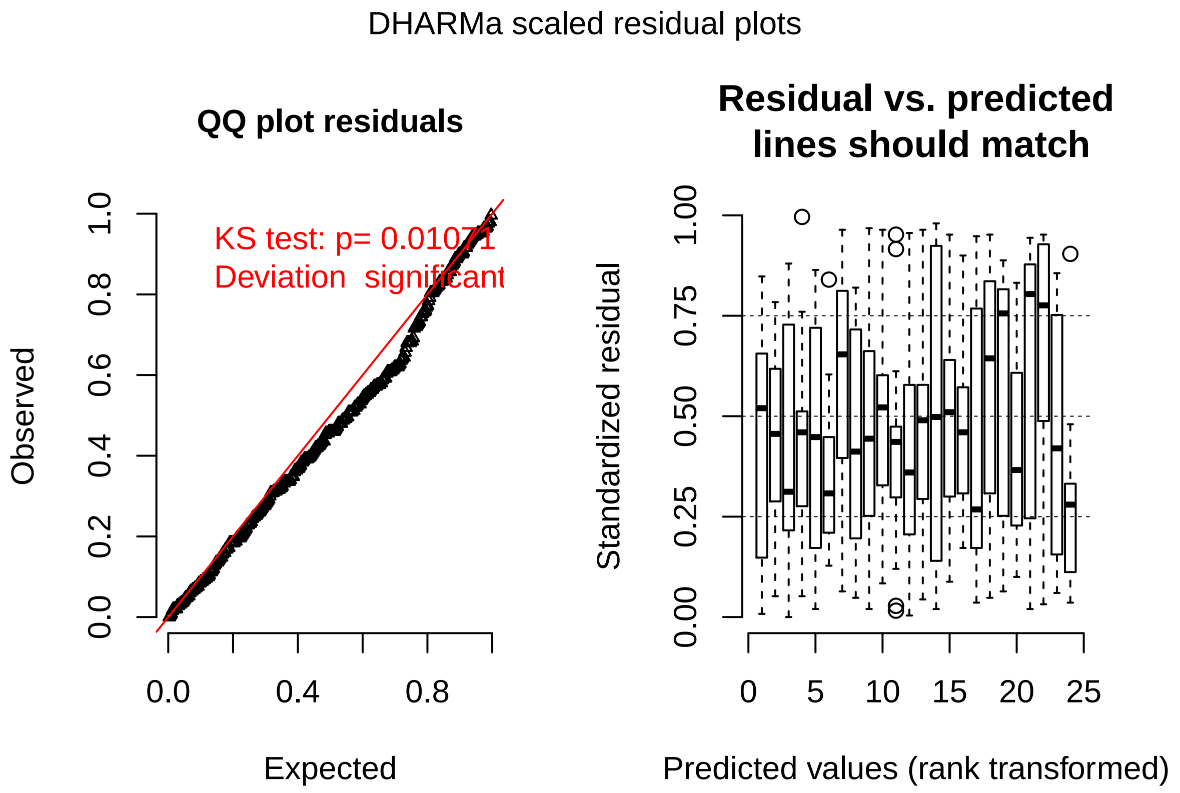

the model is zero inflated. let's remodel

```r
mpExp3 <- zeroinfl(pherbk ~ visit*mol + 1 | colony/antID, data = riskgeo)
```

```
## Error in optim(fn = loglikfun, gr = gradfun, par = c(start$count, start$zero, : valore non finito fo
```

does not work. I will remove colony from random effect.

```
mpExp3 <- zeroinfl(pherbk ~ visit*mol + 1 |antID, data = riskgeo)

Anova(mpExp3)
```

```
## Analysis of Deviance Table (Type II tests)
##
## Response: pherbk
##           Df   Chisq Pr(>Chisq)
## visit      1  0.3297     0.5658
## mol        2 47.0827  5.972e-11 ***
## visit:mol  2  0.8738     0.6460
## ---
## Signif. codes:  0 '***' 0.001 '**' 0.01 '*' 0.05 '.' 0.1 ' ' 1
```

```
meanobj<-emmeans(mpExp3,~visit*mol,type="response")
contrast(meanobj,list(mol0.1vs0.3=c(1,-1,0),
                      mol0.1vs0.9=c(1,0,-1),
                      mol0.3vs0.9=c(0,1,-1),
                      SafeVsRisky=c(-0.5,1,-0.5)),
         adjust="bonferroni")
```

```
##  contrast      estimate        SE  df z.ratio p.value
##  mol0.1vs0.3 -0.8820426 0.1435678 Inf  -6.144  <.0001
##  mol0.1vs0.9 -1.4791219 0.1805333 Inf  -8.193  <.0001
##  mol0.3vs0.9 -0.5970792 0.1651513 Inf  -3.615  0.0012
##  SafeVsRisky  0.1424817 0.1256792 Inf   1.134  1.0000
##
## Results are averaged over the levels of: antID
## P value adjustment: bonferroni method for 4 tests
```

```
phb3<-ggplot(riskgeop,aes(x=visit,y=pherbk, color=mol))+
  labs(title = "F")+
  scale_color_brewer(name="molarity", palette="Dark2")+
  ylab("Pheromone deposited back to the nest")+
  scale_x_discrete(limits=c(1,2,3,4,5,6,7,8))+
  ylim(0,20)+
  theme_light()+
  theme(axis.text.x = element_text(size=12,colour="black"),
        axis.text.y = element_text(size=12,colour="black"),
        axis.title.x = element_text(size=14),
        axis.title.y = element_blank(),
        plot.title = element_text(size=18),
        legend.position="bottom")+
  geom_jitter(width = 0.2,height=0)+
  geom_smooth(method="loess")
```

###### 2.2.4 graph together

now I will plot the pheromone deposition all together for the three experiments

```
# Multiple plot function
#
# from: http://www.cookbook-r.com/Graphs/Multiple_graphs_on_one_page_(ggplot2)/
```

```r
#
# ggplot objects can be passed in ..., or to plotlist (as a list of ggplot objects)
# - cols:   Number of columns in layout
# - layout: A matrix specifying the layout. If present, 'cols' is ignored.
#
# If the layout is something like matrix(c(1,2,3,3), nrow=2, byrow=TRUE),
# then plot 1 will go in the upper left, 2 will go in the upper right, and
# 3 will go all the way across the bottom.
#
multiplot <- function(..., plotlist=NULL, file, cols=1, layout=NULL) {
  library(grid)

  # Make a list from the ... arguments and plotlist
  plots <- c(list(...), plotlist)

  numPlots = length(plots)

  # If layout is NULL, then use 'cols' to determine layout
  if (is.null(layout)) {
    # Make the panel
    # ncol: Number of columns of plots
    # nrow: Number of rows needed, calculated from # of cols
    layout <- matrix(seq(1, cols * ceiling(numPlots/cols)),
                    ncol = cols, nrow = ceiling(numPlots/cols))
  }

 if (numPlots==1) {
    print(plots[[1]])

  } else {
    # Set up the page
    grid.newpage()
    pushViewport(viewport(layout = grid.layout(nrow(layout), ncol(layout))))

    # Make each plot, in the correct location
    for (i in 1:numPlots) {
      # Get the i,j matrix positions of the regions that contain this subplot
      matchidx <- as.data.frame(which(layout == i, arr.ind = TRUE))

      print(plots[[i]], vp = viewport(layout.pos.row = matchidx$row,
                                      layout.pos.col = matchidx$col))
    }
  }
}

multiplot(phg1,phb1,phg2,phb2,phg3,phb3,cols=3)
```

```
## `geom_smooth()` using method = 'loess'

## Warning: Removed 64 rows containing non-finite values (stat_smooth).

## Warning: Removed 64 rows containing missing values (geom_point).

## `geom_smooth()` using method = 'loess'

## Warning: Removed 1 rows containing non-finite values (stat_smooth).
```

```
## Warning: Removed 1 rows containing missing values (geom_point).
## Warning: Removed 14 rows containing missing values (geom_smooth).
## `geom_smooth()` using method = 'loess'
## Warning: Removed 69 rows containing non-finite values (stat_smooth).
## Warning: Removed 69 rows containing missing values (geom_point).
## Warning: Removed 1 rows containing missing values (geom_smooth).
## `geom_smooth()` using method = 'loess'
## Warning: Removed 18 rows containing non-finite values (stat_smooth).
## Warning: Removed 18 rows containing missing values (geom_point).
## `geom_smooth()` using method = 'loess'
## Warning: Removed 43 rows containing non-finite values (stat_smooth).
## Warning: Removed 43 rows containing missing values (geom_point).
## Warning: Removed 29 rows containing missing values (geom_smooth).
## Warning: Removed 5 rows containing non-finite values (stat_smooth).
## Warning: Removed 5 rows containing missing values (geom_point).
## Warning: Removed 16 rows containing missing values (geom_smooth).
```

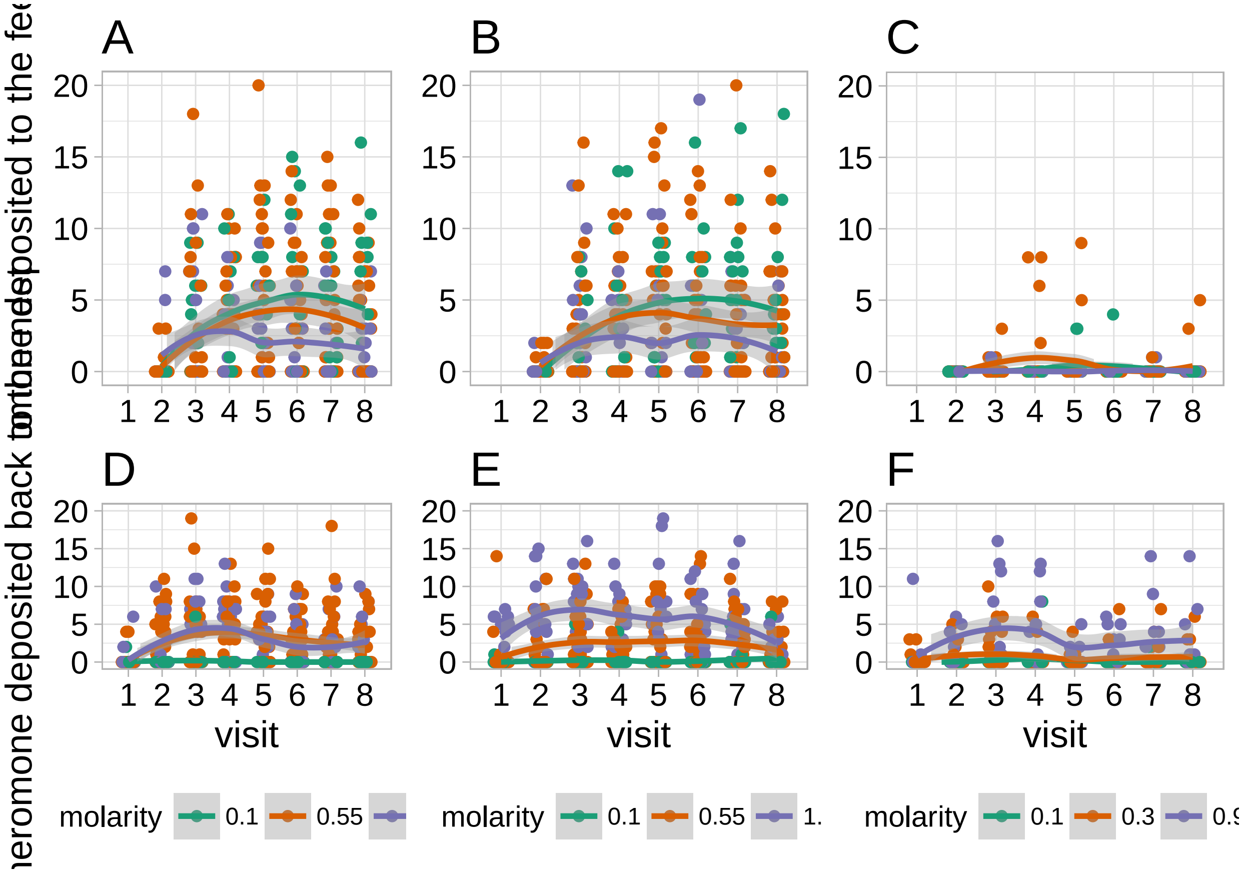

### 3 Supplemental pilot experiment

#### 3.1 Ant perception of 0.1,0.3,0.9

```
m0<-glmer(value~contrast*Visitnumber+(1|AntID),data=ctrlmelted,family="binomial",
          glmerControl(optimizer="bobyqa", optCtrl = list(maxfun = 100000000)))
Anova(m0)
```

```
## Analysis of Deviance Table (Type II Wald chisquare tests)
##
## Response: value
##                       Chisq Df Pr(>Chisq)
## contrast             3.5857  1    0.05828 .
## Visitnumber          1.4604  1    0.22686
## contrast:Visitnumber 0.0242  1    0.87635
## ---
## Signif. codes:  0 '***' 0.001 '**' 0.01 '*' 0.05 '.' 0.1 ' ' 1
```

no difference between visits

```
m1<-glmer(value~contrast*variable+(1|Colony/AntID),data=ctrlmelted,family="binomial",
          glmerControl(optimizer="bobyqa", optCtrl = list(maxfun = 100000000)))

e<-emmeans(m1,~contrast*variable,type="response")
test(e,adjust="bonferroni")
```

```
##  contrast variable          prob         SE  df z.ratio p.value
##  0.1vs0.3 Firstchoice  0.8596342 0.05688621 Inf   3.844  0.0005
##  0.3vs0.9 Firstchoice  0.6480208 0.09530360 Inf   1.461  0.5763
##  0.1vs0.3 Secondchoice 0.8699008 0.05392759 Inf   3.988  0.0003
##  0.3vs0.9 Secondchoice 0.7235134 0.08497770 Inf   2.264  0.0942
##
## P value adjustment: bonferroni method for 4 tests
## Tests are performed on the logit scale
```

#### 3.2 Discriminate three drops

```
m0<-glmer(value~Visitnumber+(1|Colony/AntID),data=melted,family="binomial",
          glmerControl(optimizer="bobyqa", optCtrl = list(maxfun = 100000000)))
Anova(m0)
```

```
## Analysis of Deviance Table (Type II Wald chisquare tests)
##
## Response: value
##              Chisq Df Pr(>Chisq)
## Visitnumber 4.5959  1    0.03205 *
## ---
## Signif. codes:  0 '***' 0.001 '**' 0.01 '*' 0.05 '.' 0.1 ' ' 1
```

difference between visits

```
m1<-glmer(value~variable+(1|Colony/AntID),data=melted,family="binomial",
          glmerControl(optimizer="bobyqa", optCtrl = list(maxfun = 100000000)))
Anova(m1)
```

```
## Analysis of Deviance Table (Type II Wald chisquare tests)
##
## Response: value
##           Chisq Df Pr(>Chisq)
## variable 0.1038  1     0.7473
```

no difference between first and last choice. I will just look at all the percentages together.

```
melted$Visitnumber<-as.factor(melted$Visitnumber)
m2<-glmer(value~variable*Visitnumber+(1|Colony/AntID),data=melted,family="binomial",
          glmerControl(optimizer="bobyqa", optCtrl = list(maxfun = 100000000)))
```

```
## Warning in checkConv(attr(opt, "derivs"), opt$par, ctrl = control$checkConv, : Model is nearly uniden
##  - Rescale variables?
```

```
e<-emmeans(m2,~variable*Visitnumber,type="response")
e
```

```
##  variable     Visitnumber      prob           SE  df    asymp.LCL
##  Firstchoice  10          1.0000000 2.484427e-07 Inf 2.220446e-16
##  Secondchoice 10          1.0000000 4.750132e-07 Inf 2.220446e-16
##  Firstchoice  11          0.9580370 5.009378e-02 Inf 6.650518e-01
##  Secondchoice 11          0.9580370 5.009390e-02 Inf 6.650502e-01
##  Firstchoice  12          0.9580370 5.009394e-02 Inf 6.650499e-01
##  Secondchoice 12          0.9580370 5.009397e-02 Inf 6.650495e-01
##  Firstchoice  13          0.9580370 5.009384e-02 Inf 6.650510e-01
##  Secondchoice 13          0.9580370 5.009396e-02 Inf 6.650496e-01
##  Firstchoice  14          0.9074061 8.321675e-02 Inf 5.844690e-01
##  Secondchoice 14          0.8481657 1.122855e-01 Inf 5.028421e-01
##  asymp.UCL
##  1.0000000
##  1.0000000
##  0.9962051
##  0.9962052
##  0.9962052
##  0.9962052
##  0.9962052
##  0.9962052
##  0.9855654
##  0.9686049
##
## Confidence level used: 0.95
## Intervals are back-transformed from the logit scale
```

I have a 100% probability of choosing safe for the first trial, the percentage decrease with subsequent, but it remains very high.

#### 3.3 fed on 1.5 risk in losses

##### 3.3.1 Preliminary questions

###### 3.3.1.1 initial vs. final

first, I want to know if initial and final choice differ

```
fsdiff<-melt(risksa15, measure.vars = c("firstchoicesafe","endchoicesafe"))
```

```r
mdiff<-glmer(value~variable+(1|colony/antID),data=fsdiff,family="binomial")
Anova(mdiff)
```

```
## Analysis of Deviance Table (Type II Wald chisquare tests)
##
## Response: value
##           Chisq Df Pr(>Chisq)
## variable 0.0952  1     0.7577
```

```r
e<-emmeans(mdiff, ~variable, type="response")
pairs(e)
```

```
##  contrast                        odds.ratio        SE  df z.ratio p.value
##  firstchoicesafe / endchoicesafe    1.09989 0.3394402 Inf   0.309  0.7577
##
## Tests are performed on the log odds ratio scale
```

there is no difference between primary and secondary choice, I will now on only use the primary for further analysis

###### 3.3.1.2 vistits n.

now, I want to know if the visits differ from one another

```r
risksa15$visit<-as.numeric(risksa15$visit)
mvisdiff<-glmer(firstchoicesafe~visit+(1|colony/antID),data=risksa15,family="binomial",
                glmerControl(optimizer="bobyqa", optCtrl = list(maxfun = 1000000000)))

Anova(mvisdiff)
```

```
## Analysis of Deviance Table (Type II Wald chisquare tests)
##
## Response: firstchoicesafe
##        Chisq Df Pr(>Chisq)
## visit 4.6318  1    0.03139 *
## ---
## Signif. codes:  0 '***' 0.001 '**' 0.01 '*' 0.05 '.' 0.1 ' ' 1
```

```r
summary(mvisdiff)
```

```
## Generalized linear mixed model fit by maximum likelihood (Laplace
##   Approximation) [glmerMod]
##  Family: binomial  ( logit )
## Formula: firstchoicesafe ~ visit + (1 | colony/antID)
##    Data: risksa15
## Control:
## glmerControl(optimizer = "bobyqa", optCtrl = list(maxfun = 1e+09))
##
##      AIC      BIC   logLik deviance df.resid
##    193.9    206.9    -93.0    185.9      185
##
## Scaled residuals:
##     Min      1Q  Median      3Q     Max
## -2.4228  0.2194  0.2921  0.3888  0.8847
##
## Random effects:
```

```
##  Groups       Name        Variance Std.Dev.
##  antID:colony (Intercept) 2.385    1.544
##  colony       (Intercept) 0.000    0.000
## Number of obs: 189, groups:  antID:colony, 63; colony, 4
##
## Fixed effects:
##             Estimate Std. Error z value Pr(>|z|)
## (Intercept)   7.5727     2.7825   2.722   0.0065 **
## visit        -0.5722     0.2659  -2.152   0.0314 *
## ---
## Signif. codes:  0 '***' 0.001 '**' 0.01 '*' 0.05 '.' 0.1 ' ' 1
##
## Correlation of Fixed Effects:
##       (Intr)
## visit -0.991
```

the percentage of ants going for safe decreases with successive visits. this means that more and more ants after not finding the sugar drop start doing a random search. I will from now on only observe the first visit, being it a clearer indication of choice

##### 3.3.2 modeling

now to the actual model. I drop antID because I kept only one observation for each ant

```
risksa15sing<-subset(risksa15,risksa15$visit==9)

mExp1<-glmer(firstchoicesafe~firstfeed*firstrisk+(1|antID),data=risksa15sing,family="binomial",
             glmerControl(optimizer="bobyqa", optCtrl = list(maxfun = 1000000)))
```

```
## Warning in checkConv(attr(opt, "derivs"), opt$par, ctrl = control$checkConv, : Model is nearly unide
##  - Rescale variables?
```

```
simres<-simulateResiduals(mExp1) #standard seed for random values is 123
plot(simres, asFactor=T)
```

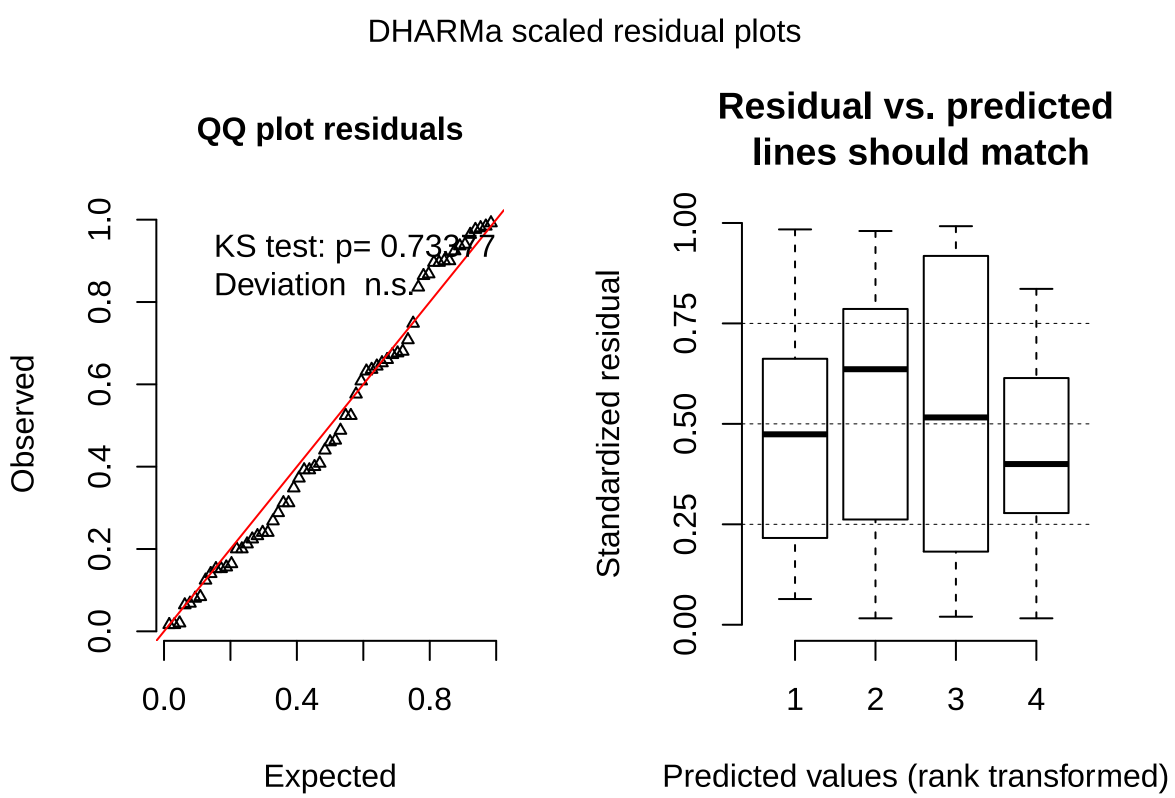

model is good here. it says nearly unidentifiable. probably I have complete separation of one data point, like 100% prob for one group. let's go on

```
Anova(mExp1)
```

```
## Analysis of Deviance Table (Type II Wald chisquare tests)
##
## Response: firstchoicesafe
##                       Chisq Df Pr(>Chisq)
## firstfeed            1.1779  1     0.2778
## firstrisk            0.9249  1     0.3362
## firstfeed:firstrisk 0.0010  1     0.9752
```

no effect of any of the factors, so I will redo the model without factors

```
mExp1<-glmer(firstchoicesafe~+(1|antID),data=risksa15sing,family="binomial",
             glmerControl(optimizer="bobyqa", optCtrl = list(maxfun = 1000000)))

simres<-simulateResiduals(mExp1) #standard seed for random values is 123
plot(simres, asFactor=T)
```

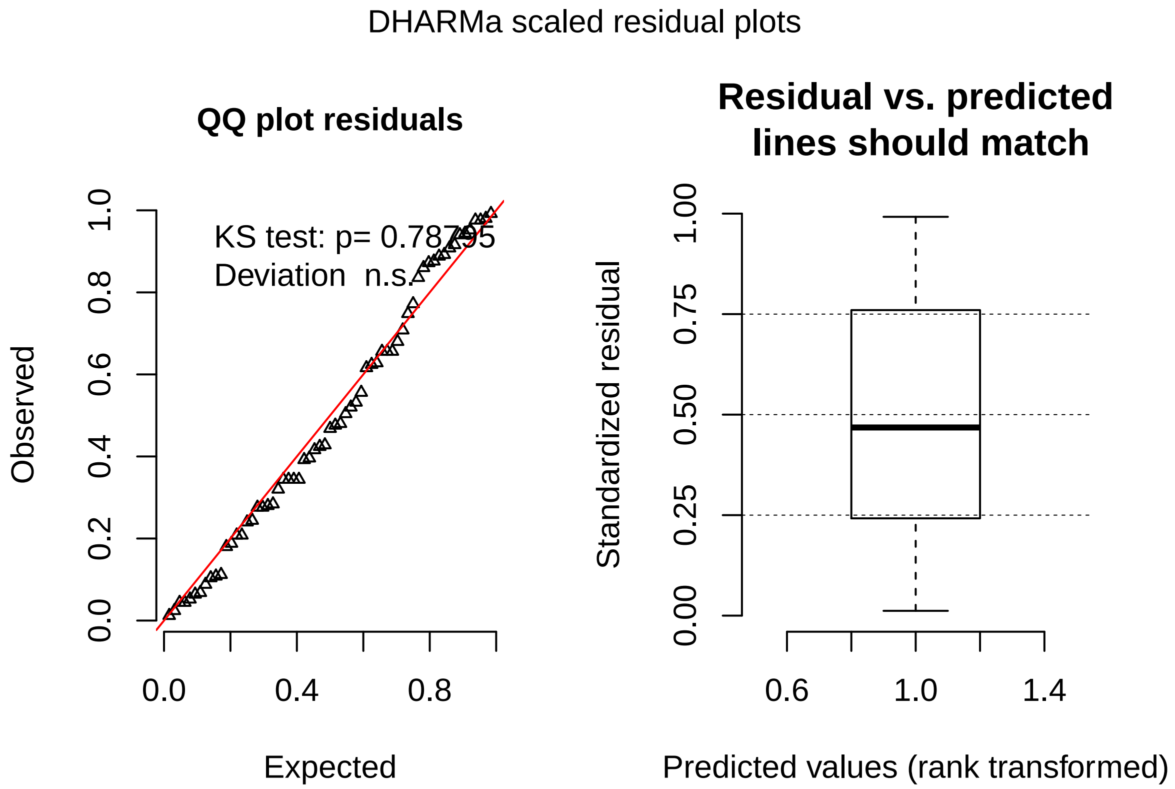

```r
meanobj <- emmeans(mExp1,~1, type="response")
kable(test(meanobj))
```

| 1 | prob | SE | df | z.ratio | p.value |
|---|---|---|---|---|---|
| overall | 0.8253968 | 0.0479891 | Inf | 4.664892 | 3.1e-06 |

ants prefer the safe **82%**.
